# A mechanistic digital twin model for epigenetic therapy optimization in triple-negative breast cancer

**DOI:** 10.64898/2026.09.11.750942

**Authors:** Simone Bruno, Alexandra Indeglia, Sophia Lichterfeld, Amy E Schade, Karen Cichowski, Franziska Michor

## Abstract

Epigenetic therapies offer a promising approach to cancer treatment by modulating chromatin states that govern tumor cell identity, plasticity and therapeutic response. However, predicting and optimizing the effects of such interventions remains challenging. Here, we developed a digital twin framework that integrates mechanistic models of chromatin regulation, *in vitro* cell-state and treatment response data, and pharmacokinetics to simulate tumor progression and therapeutic response. We applied this framework to triple-negative breast cancer (TNBC), an aggressive disease in which chromatin dysregulation contributes to tumor progression, and investigated combination treatment with an EZH2 inhibitor promiting chromatin opening and an AKT inhibitor, which together enhance expression of GATA3 and BMF. Parameterized and validated using *in vitro* treatment response data, the model enables *in silico* clinical trials of alternative combination regimens and treatment schedules. These simulations identify regimens that achieve comparable therapeutic effects to reference schedules while substantially reducing cumulative drug exposure. We further demonstrated the digitan twin’s ability of identifying personalized therapeutic strategies by incorporating patient-specific treatment-response data. Our work establishes a mechanistic digital twin framework for predicting tumor responses to chromatin-modifying therapies and provides a quantitative approach for optimizing treatment combinations and schedules across diverse cancer contexts.

## Introduction

Epigenetic therapies have emerged as a promising strategy in cancer treatment because chromatin modifications regulate gene expression and thereby control tumor cell identity, plasticity, and therapeutic response [1–3]. Increasing evidence suggests that targeting epigenetic regulators can reprogram tumor cell states and enhance the efficacy of other anticancer therapies. As a result, inhibitors of chromatin-modifying enzymes, including histone methyltransferases, histone deacetylases, and DNA methyltransferases, are increasingly being explored as components of combination therapies across multiple cancer types. One cancer type in which such approaches are being actively investigated is triple-negative breast cancer (TNBC) [4].

TNBC is the most aggressive subtype of breast cancer, with a high rate of recurrence and approximately 52,000 new cases diagnosed annually in the United States [5–7]. Current standard-of-care treatment relies on systemic chemotherapy, often combined with immunotherapy, yet these approaches frequently produce only short-lived responses [7, 8]. These limitations underscore the urgent need for more effective therapeutic strategies in TNBC. The PI3K/AKT pathway represents a plausible therapeutic target, as it is frequently hyperactivated in TNBC, with alterations reported in more than 70% of cases [9–12]. Despite this rationale, a recent Phase III clinical trial combining an AKT inhibitor with the chemotherapeutic agent paclitaxel failed to improve outcomes in TNBC patients [13, 14]. Although additional studies evaluating alternative agents and combinations are ongoing [15], the therapeutic potential of PI3K/AKT pathway inhibition in TNBC remains uncertain.

A recent study suggests a potential path forward: while PI3K pathway inhibitors have efficacy in luminal-like breast tumors [16], most TNBCs exhibit a basal-like phenotype [17], raising the possibility that altering cell state could sensitize TNBC to AKT pathway inhibition. Chromatin modifications, including histone modifications and DNA methylation, are central to the maintenance of cell state identity [18–21], and emerging experimental evidence implicates these processes in TNBC tumorigenesis and progression [22]. Consistent with this hypothesis, AKT inhibitors, when combined with inhibitors of the epigenetic regulator EZH2, decimate TNBC cells and induce marked tumor regression in multiple *in vivo* models [4]. These findings align with a growing body of preclinical and clinical evidence suggesting targeted inhibition of epigenetic modifiers as a strategy to enhance cancer therapy [23]. These results suggest that modulating chromatin state may sensitize TNBC to targeted therapies, but optimal strategies for clinical testing remain poorly defined.

Computational modeling provides a powerful approach to address these challenges by integrating mechanistic knowledge with experimental data to predict therapeutic responses and explore alternative treatment strategies [24]. However, these approaches typically represent treatment effects through pharmacological, tumor growth, immune response, or resistance models [25–27] and generally do not explicitly describe chromatin-regulatory dynamics and their impact on cell-state transitions. Even when epigenetic therapies are considered, their effects are often incorporated phenomenologically [28] rather than through a mechanistic representation of chromatin regulation. As a result, their ability to mechanistically investigate epigenetic therapies remains limited.

Recent advances in computational modeling have enabled the development of digital twins, patient-specific computational representations that integrate biological knowledge with experimental or clinical data to predict disease progression and therapeutic response [29–31]. In oncology, these approaches are increasingly being explored to predict individual treatment response, optimize therapeutic strategies, and potentially support treatment selection [29, 32, 33]. Recent applications span multiple cancer types and therapeutic modalities, including patient-specific radiotherapy optimization in glioma [34], tumor-growth prediction in prostate cancer [35], and treatment decision support in multiple myeloma [36]. Digital twins have also recently been developed for TNBC, integrating patient-specific imaging data with tumor-growth models to predict optimal therapeutic regimens [37]. However, none of these models have focused on epigenetic therapies or explicitly incorporated the chromatin-regulatory mechanisms underlying their effects. Thus, the application of digital twins to the mechanistic investigation and optimization of epigenetic therapies remains largely unexplored.

Here, we developed a digital twin model that integrates mechanistic modeling of epigenetically driven tumour dynamics with pharmacokinetics to quantitatively predict cancer progression and therapeutic response. Calibrated and validated using experimental data, the framework enables *in silico* evaluation and optimization of epigenetic combination therapies, including assessment of treatment robustness and resistance evolution. We applied the model to triple-negative breast cancer (TNBC) to identify a clinically feasible treatment schedule that optimizes outcomes across a patient cohort and, using patient-specific response data, to derive personalized optimal schedules. This framework provides a rational approach for designing and optimizing epigenetic therapies and can be adapted to other chromatin modification-driven cancers.

## Results

### Development of the digital twin model

Our digital twin model integrates three components: a mechanistic mathematical model of epigenetically driven tumor progression dynamics, *in vitro* experimental data measuring the response of tumor cells to epigenetic combination therapy, and pharmacokinetic (PK) models predicting the concentration of each drug over time in patients (Fig. 1A). The tumor progression model is calibrated and validated using experimental measurements and then coupled with PK model-derived drug concentration profiles to predict therapeutic responses to different combination dosing regimens. The resulting digital twin can quantify outcomes such as cell death for individual combination strategies and identify optimal treatment schedules. Depending on the available data, the same framework can be used either to identify treatment schedules that perform robustly across heterogeneous patient cohorts or to identify personalized optimal treatment schedules using patientspecific treatment response data. The digital twin model can be readily adapted to different cancer types and epigenetic therapies. As a demonstration of its capabilities, we applied the framework to the design and optimization of epigenetic therapies in TNBC.

**Figure 1.**
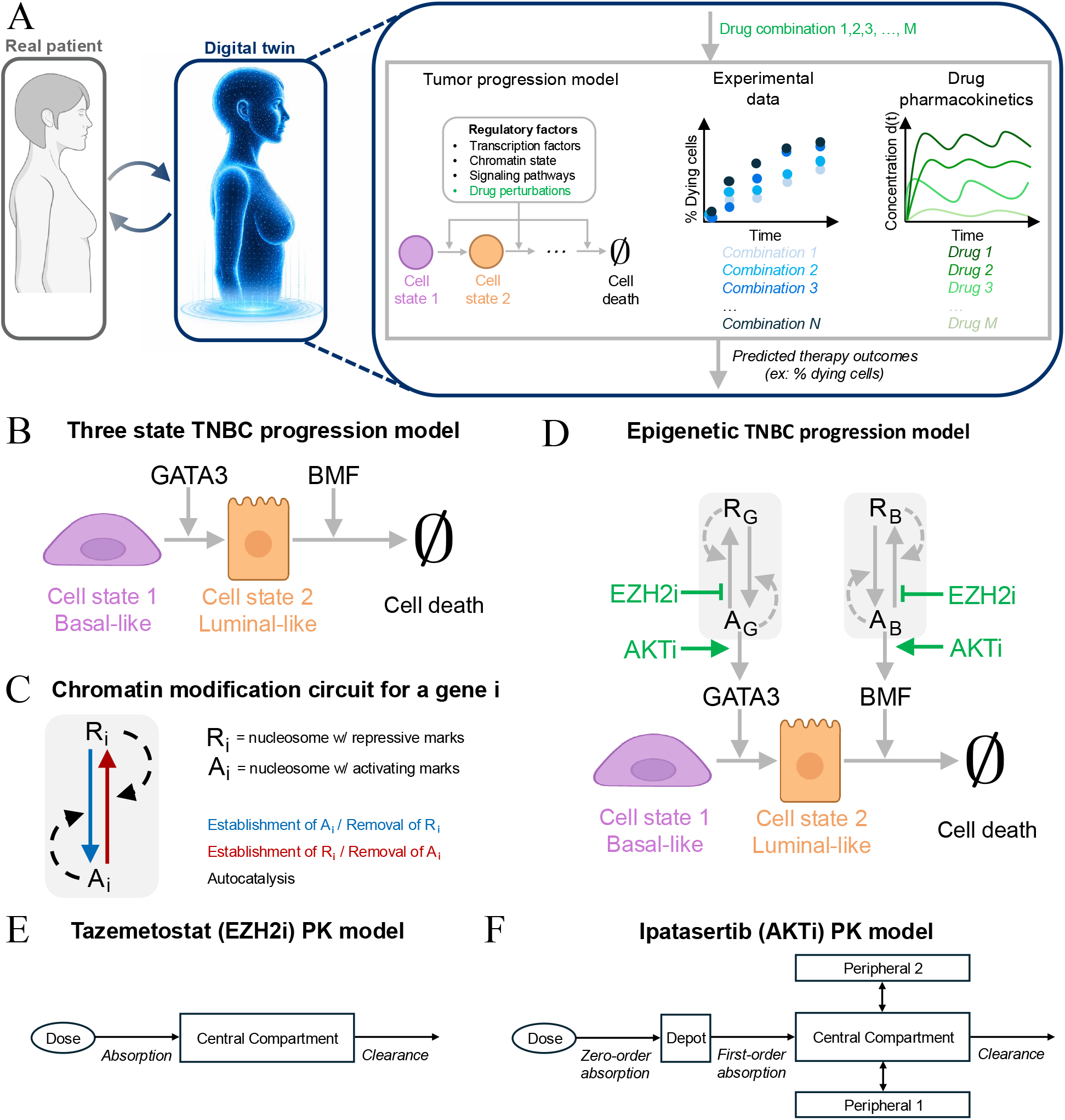
A digital twin model for epigenetic therapy optimization combining mechanistic modeling, data-driven inference, and drug pharmacokinetics. **A** Schematic of the digital twin combining the tumor progression model, inference from experimental data, and PK models to investigate treatment outcomes under different drug regimens. **B** Diagram of the TNBC progression model. Basal-like TNBC cells can transition to a luminal-like state and subsequently undergo cell death, regulated by GATA3 and BMF. **C** Chromatin modification circuit for GATA3 and BMF, where nucleosomes carry either activating (A_i_, e.g., H3K4me3) or repressive (R_i_, e.g., H3K27me3) marks. The circuit includes *de novo* establishment, autocatalysis, and erasure processes. **D** The epigenetic TNBC progression model during treatment with epigenetic modifiers (drug action shown in green). **E** Pharmacokinetic (PK) model for tazemetostat (EZH2i) [38]. **F** Pharmacokinetic (PK) model for ipatasertib (AKTi) [39].

To investigate how chromatin modifications, and drugs affecting them, drive TNBC evolution, we first developed an epigenetic TNBC progression model, a mathematical framework describing the transitions between key cellular states in TNBC (Fig. 1B). In the model, cells can exist in one of three states: basal-like, luminal-like, or cell death, as previously observed [4]. TNBC cells, which are predominantly basal-like, can be reprogrammed toward a luminal-like phenotype due to the combined effect of EZH2 and AKT inhibitors, and the luminal-like state can subsequently transition to apoptosis. This sequence encapsulates the therapeutic rationale that modulating TNBC cells from a basal-like to a luminal-like state, typical of other breast cancer subtypes in which AKT inhibitors have already been shown to be effective [40], sensitizes TNBCs to AKT inhibitors.

Two molecular regulators have been shown to be critical for these transitions [4]. GATA3, a master transcription factor controlling luminal differentiation, is upregulated only upon combined EZH2 and AKT inhibition, establishing its role in mediating the basal-to-luminal transition. BMF, a proapoptotic BH3-only protein, is the most strongly induced apoptotic gene under the same combined treatment, and its expression is required to drive the subsequent transition from the luminal-like state to apoptosis. Together, these findings establish GATA3 and BMF as key regulators of TNBC cell-state transitions (Fig. 1B).

To mechanistically capture how chromatin modifications regulate these processes, we introduced a chromatin modification circuit for the GATA3 and BMF genes (Fig. 1C). This circuit is built on previous models of chromatin modification dynamics [21, 41, 42] based on detailed characterizations of these modifications [43–47]. In this model, for each gene i of interest, each nucleosome can be modified either with a repressive chromatin mark, such as H3K27me3 (R_i_), or with an activating mark, such as H3K4me3 (A_i_). H3K4 methylation is a histone modification that promotes a less compact DNA structure and is therefore associated with an active chromatin state [18, 45], whereas H3K27 methylation compacts the DNA around nucleosomes and are thus associated with a repressed chromatin state [20].

In the model, activating chromatin marks are established *de novo* by writer enzymes and reinforce their own propagation through autocatalytic recruitment of these writers (Fig. 1C). Repressive marks are introduced and maintained through an analogous autocatalytic process. Both classes of modifications are passively removed through dilution during DNA replication or basal demethylation or deacetylation [18] (Fig. 1C). In addition, activating and repressive marks promote each other’s removal by recruiting eraser enzymes for the opposing modification (Fig. 1C). To simplify the system, we assume that each nucleosome carries either an activating or a repressive modification, with no unmodified state. This reduction in state space limits the number of parameters while preserving the autocatalytic feedback loops that underlie histone modification maintenance [21].

The model incorporates the action of two therapeutic agents: an EZH2 inhibitor (EZH2i) and an AKT inhibitor (AKTi). EZH2 is the catalytic subunit of PRC2, which deposits the repressive histone mark H3K27me3 [48] (Fig. 1D). EZH2 inhibition reduces H3K27me3 levels, thereby shifting chromatin toward a transcriptionally active state. AKT is a central signaling kinase that represses GATA3 transcription by phosphorylating and inhibiting FOXO1, preventing FOXO1-mediated gene activation [4]. AKT inhibition relieves this repression, enabling FOXO1-driven GATA3 expression. In addition, AKTi activates the STING–TBK1 pathway, which induces the IL-6–JAK1–STAT3 signaling cascade and leads to robust upregulation of the pro-apoptotic gene BMF [4]. Through these mechanisms, AKTi promotes expression of both GATA3 and BMF (Fig. 1D).

By integrating cell-state transitions with gene-level chromatin modification dynamics, we obtain the ordinary differential equation (ODE)-based epigenetic TNBC progression model (SI-Section S.1 and SI-Eq. (S.1)). This mechanistic model enables quantitative prediction of how chromatin modifications affect TNBC progression and treatment response, describing how combinatorial therapies can reprogram basal-like TNBC cells into luminal-like states and ultimately induce apoptosis.

We first parameterized the epigenetic TNBC progression model using cell culture experiments providing *in vitro* data that quantified TNBC cell response under control, single-drug, and combination treatments (see next section). These data were used to calibrate the model and identify optimal treatment schedules that perform robustly across heterogeneous patient cohorts. The same calibration framework can then be used with patient-specific treatment response data to personalize the digital twin and optimize treatment schedules for individual patients; as a proof of concept, we demonstrate this approach using synthetic patient-specific data. To translate treatment schedules into time-dependent drug exposure, we incorporated pharmacokinetic (PK) models describing inhibitors targeting EZH2 and AKT. As an illustrative example, we considered the EZH2 inhibitor tazemetostat and the AKT inhibitor ipatasertib, for which PK models have been previously reported. Tazemetostat is an FDA-approved EZH2 inhibitor [38], and based on published pharmacokinetic data [38, 49, 50], we derived a one-compartment model with first-order absorption that accurately reproduces its reported PK profile [38] (Fig. 1E and SI-Section S.3). Ipatasertib is a selective ATP-competitive AKT inhibitor currently in late-stage clinical development for the treatment of solid tumors, including breast cancer [39]. PK data from multiple phase 1 and phase 2 studies were analyzed [39], and a three-compartment model with first-order elimination and sequential zero- and first-order absorption was determined to best describe its pharmacokinetics (Fig. 1F and SI-Section S.4).

### Parameter estimation and validation using *in vitro* data

To estimate the parameters of the TNBC progression model (Fig. 1D and SI-Eq. (S.1)), we performed *in vitro* IncuCyte live-cell imaging experiments quantifying the response of TNBC cells under different treatment conditions, including untreated control and single or combined administration of the EZH2 inhibitor tazemetostat and the AKT inhibitor ipatasertib (Fig. 2A). These experiments provided longitudinal measurements of treatment-induced cell death for each condition considered (Fig. 2B), corresponding to the apoptotic state in our model (Methods). These data were then analyzed with a Bayesian inference framework, which yields posterior distributions for all parameters, quantifying uncertainty arising from measurement noise and model stochasticity (Methods).

**Figure 2.**
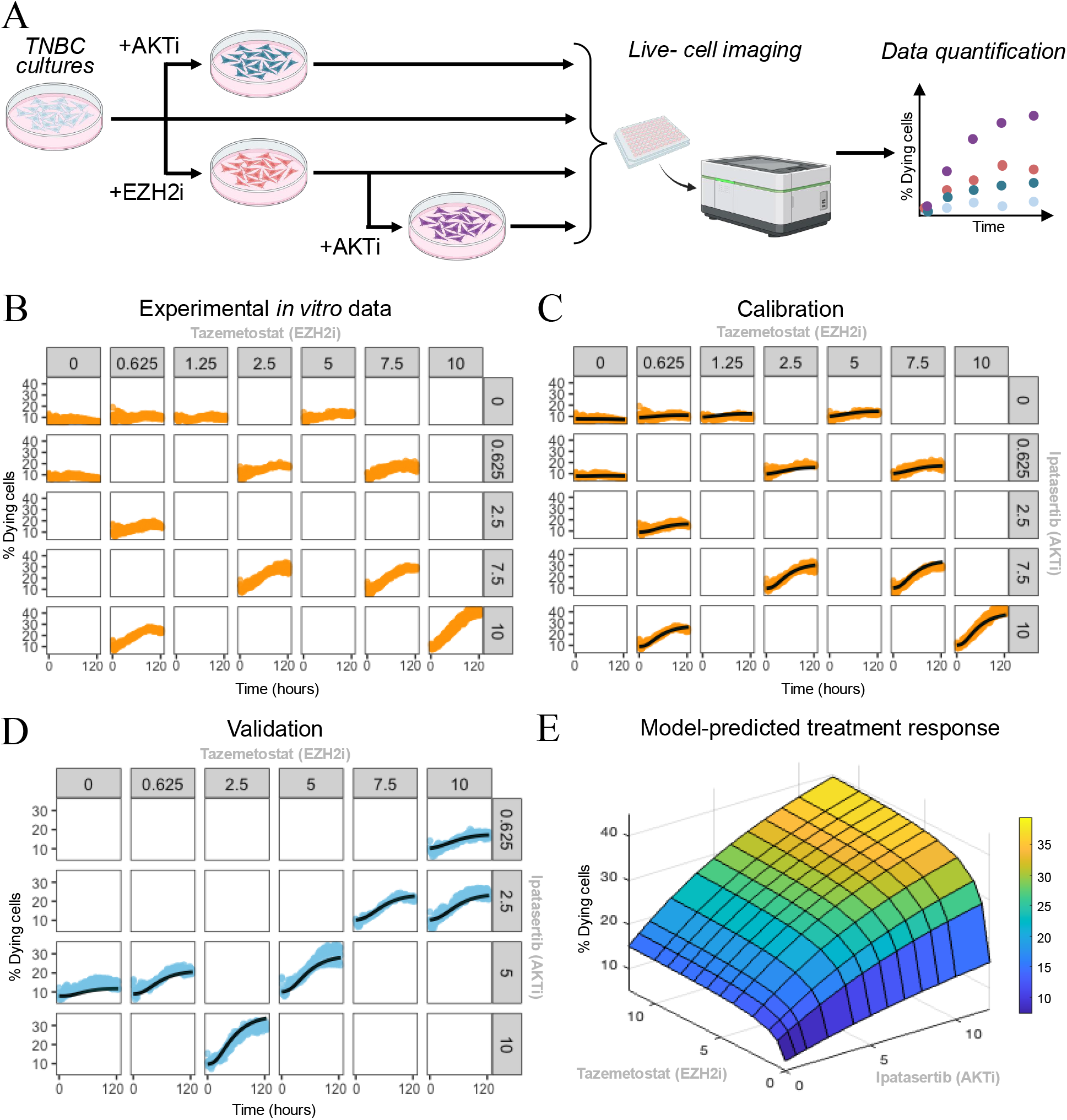
*In vitro* TNBC treatment response data and model-based predictions under AKT and EZH2 inhibition. **A** Schematic representation of the *in vitro* experiments used to generate the data for calibrating and validating the TNBC progression model. TNBC cultures were treated with AKTi, EZH2i, or their combination, followed by live-cell imaging and quantification of the percentage of dying cells over time for each condition. **B** Data used for the model parameterization, which provides longitudinal measurements of cell death across multiple drug-combination conditions. The concentrations of the AKTi and EZH2i are displayed in *µ*M. **C** Model fit to the training data. Orange denotes the experimentally determined trajectories of dying cells, and black represents the posterior mean trajectory predicted by the calibrated model. **D** Model validation. Blue shows experimentally determined trajectories of dying cells during conditions not used during parameter estimation, and black represents the corresponding posterior predictive mean obtained from the calibrated model. **E** Model-predicted three-dimensional response surface obtained by simulating the calibrated TNBC progression model across a range of ipatasertib (x-axis) and tazemetostat (y-axis) concentrations. Treatment efficacy was quantified as the percentage of dying cells at 120 hours post treatment initiation (z-axis).

The calibrated model accurately recapitulates the experimental trajectories used to estimate its parameters (Fig. 2C), with strong agreement between posterior predictive means and experimental mean trajectories (Pearson *r* = 0.97, normalized RMSE = 1.1), demonstrating that its mechanistic structure properly captures the dynamics of treatment-induced TNBC cell death.

In order to validate our model and assess its generalizability, we evaluated its predictions on drugcombination experiments not used during parameter estimation. The model is able to replicate these treatment response trajectories (Fig. 2D), indicating that the calibrated model maintains predictive accuracy beyond the conditions used for parameter estimation (Pearson *r* = 0.97 and normalized RMSE = 1.2).

### Distinct mechanisms of AKTi and EZH2i shape TNBC treatment response

The validated TNBC progression model was then used to quantitatively characterize how different concentrations of AKTi and EZH2i affect treatment response under constant exposure. Treatment efficacy was quantified as the percentage of dying cells at 120 hours. The model-predicted combination dose–response surface showed that neither AKTi nor EZH2i alone produced substantial cell death, indicating very low treatment efficacy when the drugs are used individually. By contrast, their combination produced a marked increase in the percentage of dying cells (Fig. 2E), consistent with the experimental observations (Fig. 2B,D).

Beyond reproducing these qualitative trends, the model enables quantitative comparison of dose–response behaviors across the full concentration range, revealing distinct regimes for the two inhibitors. Increasing AKTi concentration produced a gradual and sustained improvement in treatment efficacy, while increasing EZH2i concentration improved efficacy but quickly reached a plateau (Fig. 2E). This difference reflects their distinct mechanisms of action (Fig. 1D): EZH2i promotes chromatin opening, and once chromatin accessibility is maximized, further increases in concentration cannot enhance the transition to the luminal and dying states. In contrast, AKTi directly regulates GATA3 and BMF expression, allowing its effect to persist across a broader concentration range. Overall, the model enables quantitative distinction between regimes of early saturation and sustained improvement in efficacy for different inhibitors. These differences cannot be directly inferred from pathway knowledge alone but instead arise from the integration of experimental data with chromatin remodeling dynamics, transcriptional regulation, and cell-state transition modeling.

Furthermore, the model predicted that the most effective *in vitro* regimens were those combining tazemetostat concentrations of at least 5 µM with the highest tested doses of ipatasertib (Fig. 2E), and these predictions matched the experimental observations (Fig. 2C). Together, these results validate the predictive accuracy of the calibrated model and support its use in designing and evaluating treatment schedules.

### *In silico* clinical trials identify optimal treatment schedules

To identify optimal treatment schedules, we simulated *in silico* clinical trials comprising 500 virtual patients generated using our digital twin model (Fig. 3A). For each virtual patient, the pharmacokinetics of tazemetostat and ipatasertib were modeled as previously described (Fig. 1E,F and SI-Sections S.3,S.4), with inter-individual variability incorporated through the population PK models. Heterogeneity in pharmacodynamic response was captured by sampling model parameter values from the posterior distributions estimated using the Bayesian framework applied to the *in vitro* data (Methods). Each virtual patient was then simulated following a candidate treatment cycle schedule for 26 days, including 5 pretreatment days with tazemetostat only, matching the experimental pretreatment protocol used to generate the *in vitro* data for model parameterization.

**Figure 3.**
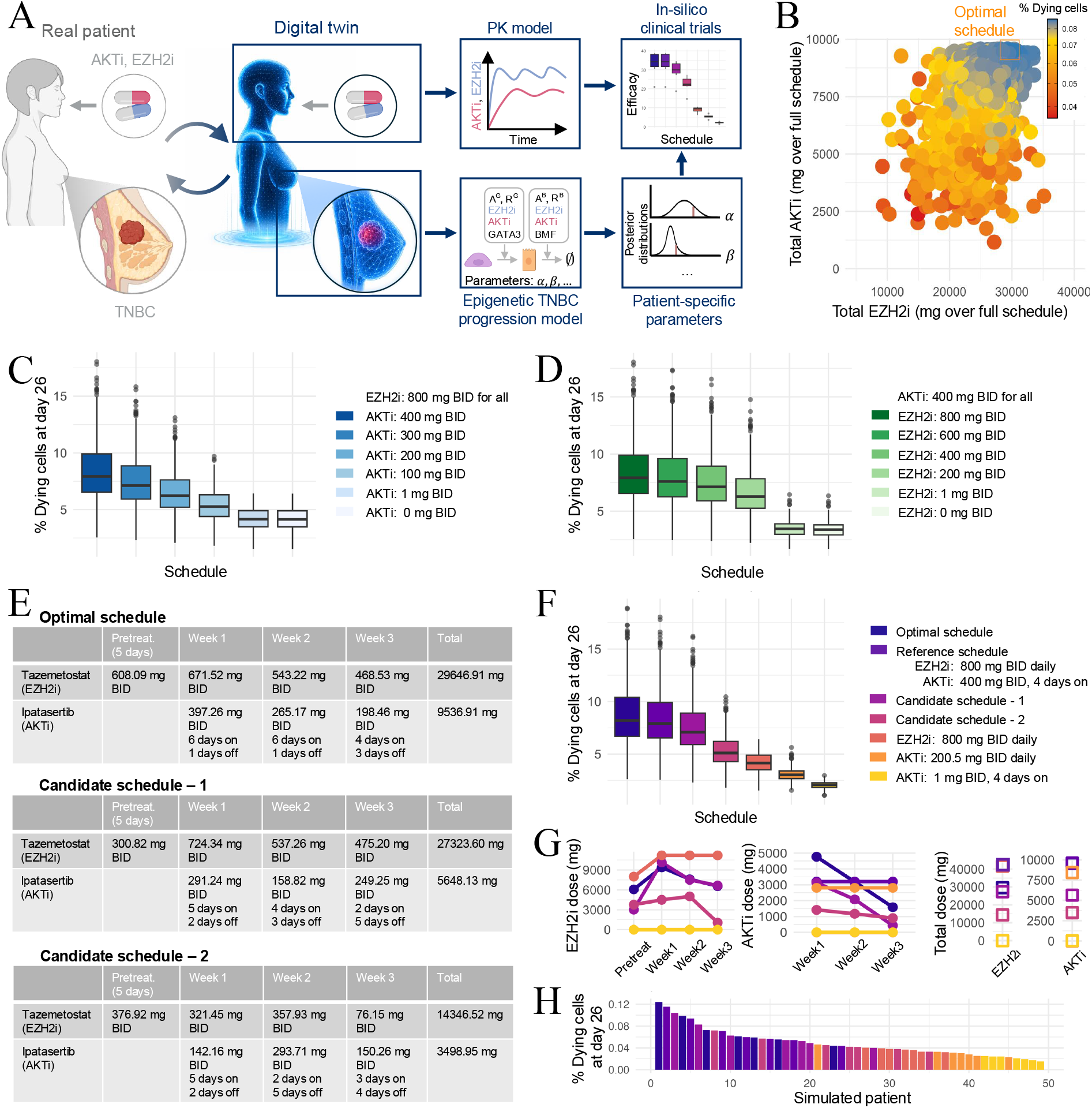
*In silico* clinical trials identify highly effective regimens with lower overall drug use. **A** Overview of the *in silico* clinical trial framework. **B** The genetic algorithm (GA) evaluates N = 4032 treatment schedules, shown according to their total tazemetostat and ipatasertib exposures across the entire treatment duration. The orange square marks the schedule identified by the GA as optimal and the general structure of candidate schedules is described in SI-Fig. S.1. **C** Box plots of regimen efficacy, expressed as the percentage of dying cells at day 26 (end of one treatment cycle). The schedules shown share the same EZH2i dosing regimen. **D** Same as in panel C, but for schedules sharing the same AKTi dosing regimen. **E** Summary of optimal and candidate schedules considered in panels F-H. **F** Box plots of regimen efficacy for the reference schedule, the optimal schedule identified by the GA, two additional candidate schedules, and single-drug schedules. For panels C, D, and F, boxes represent the middle 50% of the data, with the median shown as a horizontal line. Whiskers extend to the minimum and maximum values excluding outliers, which are shown as dots. Each regimen is represented with a different color and its results are based on N = 500 simulations, corresponding to 500 *in silico* patients. **G** Total drug exposure for each regimen considered in panel F. For both tazemetostat and ipatasertib, dots indicate the total dose delivered during the pre-treatment phase and each treatment week (weeks 1–3), and squares indicate the overall cumulative dose across the entire treatment cycle. **H** Waterfall plot of simulated patients ordered by decreasing efficacy, expressed as the percentage of dying cells at day 26. Each bar represents an individual patient and is depicted using the color assigned to the regimen received, using the same color scheme as in panel F. For each regimen, we considered 7 *in silico* patients.

Treatment efficacy was quantified as the percentage of dying cells at the end of the simulation. Candidate schedules were optimized using a genetic algorithm (GA) [51] (Fig. 3B, Methods), a population-based stochastic search method widely used for optimizing dosing regimens [52]. The GA explored schedules under clinically motivated constraints, with tazemetostat doses varying between 0 and 800 mg BID, administered daily, and ipatasertib doses between 0 and 400 mg BID, with administration frequencies ranging from daily doses to a single dose per week. The total dose of each drug was constrained not to exceed that of the reference schedule consisting of tazemetostat 800 mg BID daily and ipatasertib 400 mg BID administered four days per week (SI-Fig. S.1).

Using our digital twin model, we then compared the predicted efficacy of schedules with a fixed dose of tazemetostat and varying ipatasertib, and vice versa. Increasing ipatasertib produces a smooth, monotonic rise in treatment efficacy (Fig. 3C). In contrast, varying tazemetostat while keeping ipatasertib constant shows an early increase in efficacy that rapidly reaches a plateau, indicating that additional EZH2 inhibition would not further improve treatment responses (Fig. 3D). These distinct patterns mirror the different mechanisms of action of the two inhibitors captured in the progression model (Fig. 1D).

The *in silico* clinical trial analysis revealed that regimens involving only tazemetostat or ipatasertib consistently ranked among the least effective, while regimens combining both drugs substantially increased treatment efficacy (Fig. 3E,F). Additionally, optimization results demonstrate that adjusting both the weekly doses of the two drugs and the number of days on which ipatasertib is administered each week enables the identification of schedules that achieve efficacy comparable to the reference regimen (Fig. 3F) while requiring less ipatasertib and substantially less tazemetostat (Fig. 3E,G). This result is consistent with the drug-specific dynamics previously observed, where tazemetostat’s effectiveness rapidly plateaued, indicating that beyond a certain level, further increases do not enhance its effect. Importantly, these conclusions remain consistent across multiple efficacy metrics and evaluation time points, remaining consistent when treatment response was assessed using cell survival AUC (SI-Fig. S.2A), cumulative death induction (SI-Fig. S.2B), and the percentage of dying cells after two treatment cycles instead of one (SI-Fig. S.2C).

These findings are further supported by individual *in silico* patient dynamics (Fig. 3H), showing a subset of simulated patients randomly sampled from each regimen and ranked by their percentage of dying cells at day 26. Patients treated with optimal and reference regimens predominantly cluster in the higher efficacy regime, while those treated with other regimens appear progressively towards less efficacious regimes, with the single-drug regimens concentrated on the least effective areas.

Finally, because oral therapies such as tazemetostat and ipatasertib are administered as tablets with fixed dose, clinically feasible dosing regimens must consist of discrete multiples of the tablet dose. We therefore additionally evaluated discretized versions of the optimized and candidate regimens, corresponding to the closest realizable schedules under these dosing constraints (SI-Fig. S.3A). These practical schedules largely preserved the overall ranking observed for the continuous optimization (SI-Fig. S.3B,C).

### Optimal treatment schedules are robust to drug resistance

Despite initially effective treatment strategies, resistance may evolve and eventually reduce therapeutic efficacy. To assess the robustness of optimal treatment schedules to resistance, we extended the TNBC progression model to include a simple drug-resistance mechanism in which cells may enter a resistant state either from the basal-like or luminal-like state. The resistance model includes cell fate changes between these sensitive and resistant states, while the rest of the TNBC progression model remains unchanged (Fig. 4A, SI-Eq. (S.18)). The parameters of the original TNBC model are given by the means of the posterior distributions previously estimated (SI-Eq. S.10). In contrast, the parameters associated with the resistance mechanism (*a*_1_, *a*_2_, *b*_1_, *b*_2_ in Fig. 4A) were selected to represent three distinct resistance regimes defined by the balance between the rates at which cells transition into resistant states (*a*_1_, *a*_2_) and the death rates of resistant cells (*b*_1_, *b*_2_). Specifically, “no resistance” corresponds to the absence of transitions into resistant states (*a*_1_ = *a*_2_ = *b*_1_ = *b*_2_ = 0), “low resistance” corresponds to regimes in which resistant cells are generated but efficiently cleared (high *b*_1_, *b*_2_ or low *a*_1_, *a*_2_), and “high resistance” corresponds to regimes in which resistant cells accumulate due to high transition rates and low death rates (high *a*_1_, *a*_2_ and low *b*_1_, *b*_2_), leading to a substantial reduction in treatment-induced cell death (see SI—Section S.5 for detailed analysis). Our results show that, although resistance negatively affects the overall treatment response, the previously identified optimal schedule remains the best-performing option (Fig. 4B).

**Figure 4.**
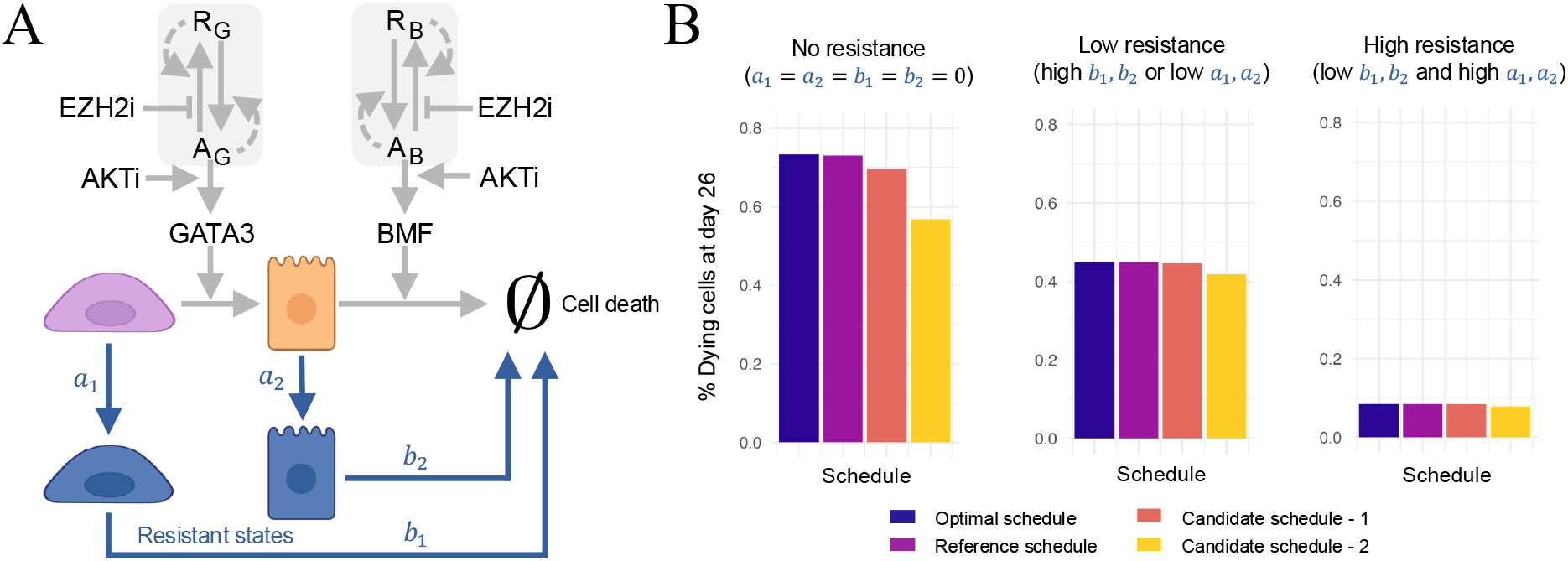
Drug resistance affects treatment efficacy. **A** Schematic diagram of a potential epigenetic TNBC progression circuit including drug-resistant states (blue). **B** % of dying cells under three resistance scenarios (no resistance, low resistance, and high resistance) obtained by simulating the ODE system of the TNBC progression model including resistant states (SI-Eq. (S.18)). Here, *a*_1_ = *a*_2_ = *b*_1_ = *b*_2_ = 0 (no resistance); *a*_1_ = *a*_2_ = 0.001 h^−1^, *b*_1_ = *b*_2_ = 0.005 h^−1^ (low resistance); and *a*_1_ = *a*_2_ = 0.01 h^−1^ *b*_1_ = *b*_2_ = 0.005 h^−1^ (high resistance). The complete list of parameter values used for these simulations is provided in SI–Section S.5, and simulations for different parameter regimes are shown in SI–Figs. S.4, S.5. The optimal schedule and candidate schedules 1 and 2 are the same as introduced in Fig. 3E.

### Impact of alternative AKT inhibitors on treatment response

Thus far, our experimental and computational analyses have focused on ipatasertib as the AKT inhibitor. Given the inherent uncertainty of drug development and the potential impact of pharmacokinetic variability on treatment efficacy, we next examined capivasertib, an alternative AKT inhibitor with comparable pharmacodynamic effects but a distinct pharmacokinetic profile [40, 53]. Substituting the ipatasertib pharmacokinetic model with that of capivasertib in our digital twin model allowed us to systematically evaluate treatment responses and identify an optimized dosing regimen for this alternative AKTi.

Capivasertib follows a three-compartment pharmacokinetic model, similar to ipatasertib, with the same overall structure for distribution and elimination. The key difference lies in the absorption process: while ipatasertib exhibits sequential absorption, with an initial zero-order phase followed by first-order absorption (Fig. 1F), capivasertib is characterized by parallel absorption, in which zeroand first-order processes occur simultaneously, with approximately 20% and 80% of the administered dose absorbed via each pathway, respectively [53] (Fig. 5A and SI-Section S.6). We considered the parameters of the TNBC progression model, estimated from experiments conducted with ipatasertib, also apply to the capivasertib treatment response, such that the two drugs have comparable PD effects and differ only in their PK properties.

**Figure 5.**
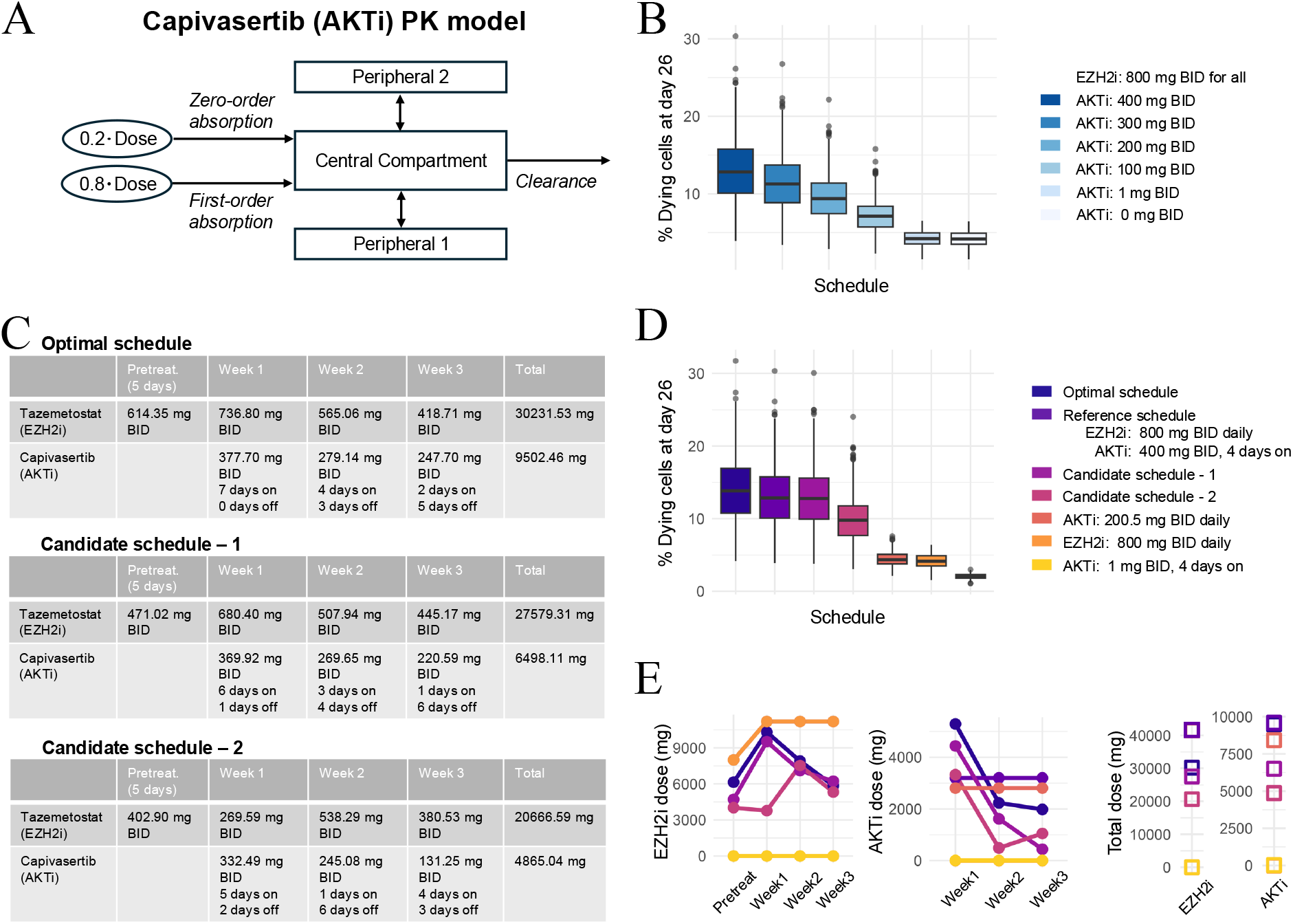
Alternative AKT inhibitors can improve treatment efficacy. **A** PK models for capivasertib (AKTi). See main text and [53] for further details. **B** Box plots of regimen efficacy, expressed as the percentage of dying cells at day 26. The schedules shown share the same EZH2i dosing regimen. **C** Summary of optimal and candidate schedules considered in panels D,E. **D** Box plots of regimen efficacy for the reference schedule, the optimal schedule identified by the Genetic Algorithm, two additional candidate schedules, and single-drug schedules. The general structure of candidate schedules is described in SI-Fig. S.1. For panels B and D, boxes represent the middle 50% of the data, with the median shown as a horizontal line. Whiskers extend to the minimum and maximum values excluding outliers, which are shown as dots. Each regimen is represented with a different color and its results are based on N = 500 simulations, corresponding to 500 *in silico* patients. **E** Total drug exposure for each regimen considered in panel D. For both tazemetostat (EZH2i) and capivasertib (AKTi), dots indicate the total dose delivered during pretreatment and each treatment week (Weeks 1–3), and squares indicate the overall cumulative dose across the entire treatment cycle.

Using our model, we first compared schedules with fixed tazemetostat and varying capivasertib (Fig. 5B). Increasing capivasertib produced a smooth and monotonic rise in treatment efficacy, similar to what was observed for ipatasertib (Fig. 3C), but the increase occurred at lower capivasertib concentrations. This shift can be explained by differences in PK properties, which allow capivasertib to reach higher drug concentrations than ipatasertib for the same administered dose.

We then conducted *in silico* clinical trials and applied our GA (Methods and SI-Fig. S.6) to identify the optimal dosing schedule (Fig. 5C-E). The optimal regimen obtained for capivasertib was structurally similar to the one identified for ipatasertib (Fig. 3E-G), exhibiting an analogous weekly pattern of AKTi administration. Specifically, in both cases the optimized schedules identified by our platform featured sustained AKTi exposure during the first treatment week, followed by a decreasing number of dosing days in subsequent weeks, suggesting that similar temporal dosing patterns arise for both inhibitors. At the same time, differences in pharmacokinetics shift the dose–efficacy tradeoff, allowing capivasertib to achieve comparable responses at lower administered doses (Fig. 5B). As a result, we can identify regimens with slightly reduced efficacy but substantially lower total drug exposure (Fig. 5C-E), highlighting clinically relevant opportunities for dose reduction without major loss of therapeutic benefit.

These conclusions remained consistent across multiple efficacy metrics, including cell survival AUC (SI-Fig. S.7A) and cumulative death induction (SI-Fig. S.7B), as well as across evaluation time points (SI-Fig. S.7C). Furthermore, because capivasertib is also administered as a fixedstrength oral tablet, we evaluated the corresponding clinically feasible discretized regimens (SI-Fig. S.8A). These discretized schedules preserved the overall ranking observed for the corresponding continuous regimens (SI-Fig. S.8B,C).

### The digital twin model can be used for individualized treatment optimization

We next sought to demonstrate, as a proof-of-concept, the use of the digital twin model for the identification of individualized optimal treatment schedules for a specific patient. To this end, we generated synthetic, patient-specific *in vitro* treatment response data (Fig. 6A and Methods) that, in practice, can be obtained from patient-derived tumor cell cultures by quantifying treatment response using our IncuCyte live-cell imaging protocol. For each virtual patient, the digital twin was then re-calibrated using these patient-specific data (Fig. 6A). Specifically, we identified *δ*_2_, *K*_*A*_, *K*_*E*_, and *δ*_1_ as the patient-specific model parameters governing treatment response (Methods and SI-Eqs. (S.1)) and re-estimated these four parameters using Bayesian inference while keeping all remaining model parameters fixed to their population-level mean values (Methods). The posterior mean values of the re-estimated parameters were then used to define the patient-specific digital twin parameters (Fig. 6B).

**Figure 6.**
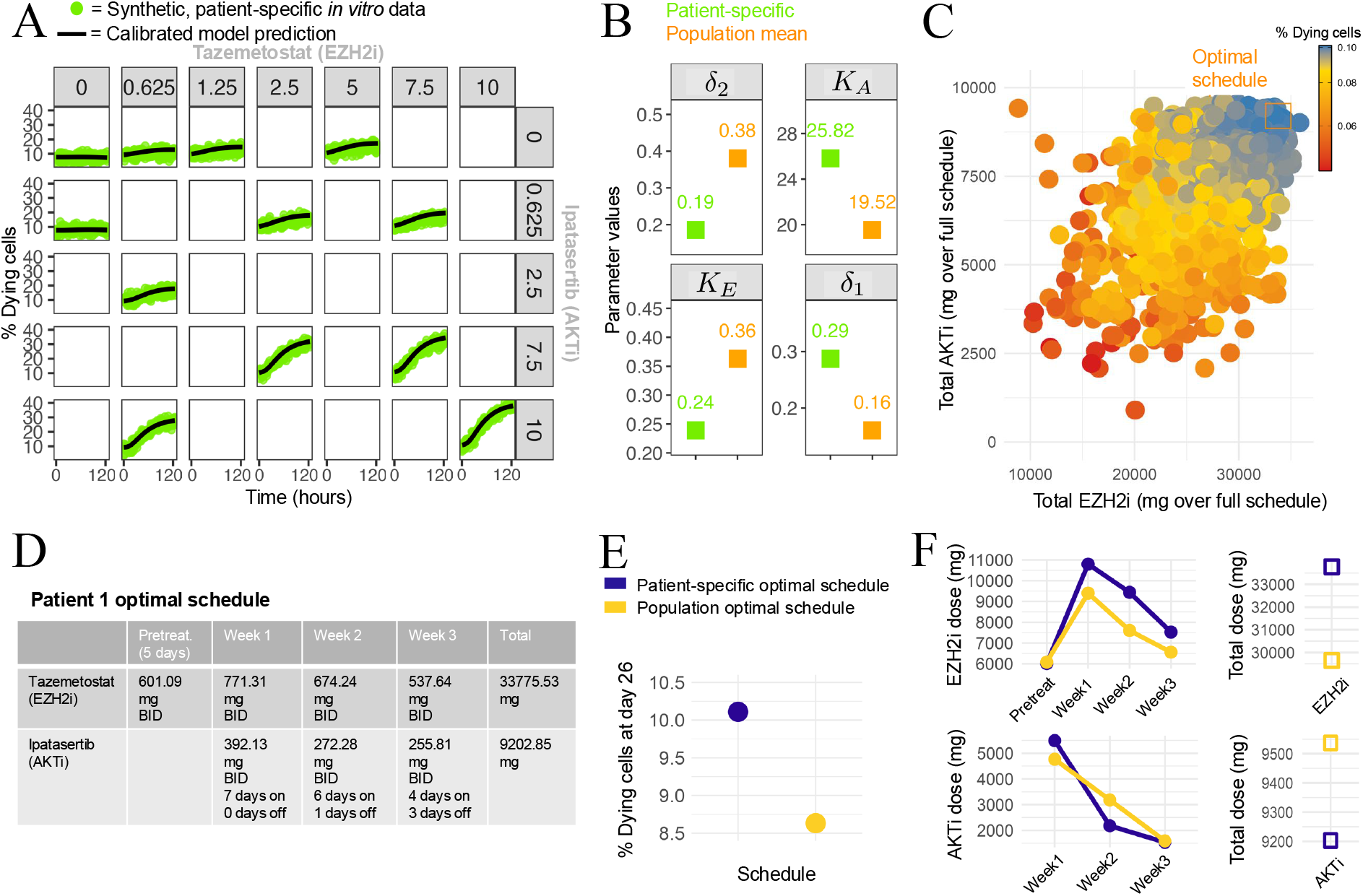
Patient-specific calibration of the digital twin model and application to individualized treatment optimization. **A** Calibration of the digital twin model using synthetic patient-specific *in vitro* treatment response data for a representative virtual patient (see Methods). Green dots denote synthetic patient-specific experimental data and black lines show the calibrated model predictions. The concentrations of the AKTi and EZH2i are displayed in *µ*M. **B** Comparison between the patient-specific parameter estimates obtained after calibration and the corresponding population-level mean parameter values, previously estimated from the population-level calibration (see SI-Section S.2), for the four patient-specific model parameters governing treatment response (*δ*_2_, *K*_*A*_, *K*_*E*_, and *δ*_1_). **C** The genetic algorithm (GA) evaluates N = 4039 treatment schedules, shown according to their total tazemetostat and ipatasertib exposures across the entire treatment duration. **D** Summary of the patient-specific optimal treatment schedule. **E** Comparison of the predicted therapeutic response obtained using the patient-specific optimal schedule and the populationlevel optimal schedule. The population-level optimal schedule is the same as the one introduced in Fig. 3E. **F** Total drug exposure for each regimen considered in panel E. For both tazemetostat and ipatasertib, dots indicate the total dose delivered during the pre-treatment phase and each treatment week (Weeks 1–3), and squares indicate the overall cumulative dose across the entire treatment cycle.

With the personalized digital twin, we then optimized candidate treatment schedules using the same GA used for the population-level analysis (Fig. 6C and Methods), identifying a patient-specific optimal treatment schedule (Fig. 6C,D). Although the optimal patient-specific schedule shared a similar overall structure to the population-level optimum (Fig. 3E), including higher and more frequent doses during the first weeks of treatment, it exhibited individualized dose adjustments reflecting the inferred patient-specific model parameters. Comparison of the patient-specific and population-level optimal regimens also revealed that patient-specific optimization further improved the predicted therapeutic response for that individual while requiring small adjustments in the dosing schedule and overall drug exposure (Fig. 6E,F).

Finally, we successfully applied the same patient-specific treatment optimization procedure to additional representative virtual patients, demonstrating the general applicability of the proposed digital twin model for individualized treatment optimization (SI-Fig. S.9).

## Discussion

Epigenetic therapies have emerged as a promising strategy in cancer treatment because chromatin modifications regulate gene expression patterns that control tumor cell identity, plasticity, and therapeutic response [1–3]. Increasing experimental evidence suggests that targeting epigenetic regulators can reprogram tumor cell states and enhance the efficacy of other anticancer therapies. However, the magnitude of these effects and how to optimally exploit them therapeutically remain unclear. To address this question, we developed a digital twin model that integrates a mechanistic model of epigenetically driven tumor progression with experimental data and pharmacokinetic models to identify optimal epigenetic treatment strategies (Fig. 1A). We illustrate the capabilities of this model by applying it to the optimization of epigenetic therapies in TNBC, a highly aggressive disease in which epigenetic modulation has been proposed as a strategy to sensitize tumors to targeted therapies. The model couples a chromatin modification circuit for GATA3 and BMF with a progression model describing transitions from basal-like to luminal-like states and ultimately to cell death (Fig. 1D). Model parameters were inferred from *in vitro* live-cell imaging data and validated on independent drug combination experiments performed under treatment conditions distinct from those used for parameter estimation, providing support for the reliability of the model predictions (Fig. 2B-D).

By considering two representative inhibitors targeting key pathways in the model, namely the EZH2 inhibitor tazemetostat and the AKT inhibitor ipatasertib, and integrating their pharmacokinetic profiles, our digital twin model enabled the simulation of *in silico* clinical trials. These simulations allowed us to identify optimized weekly dosing schedules that improved efficacy relative to monotherapies and performed comparably to the reference schedule recommended by clinicians, while requiring substantially less overall drug exposure (Fig. 3E–H). Further analysis allowed us to evaluate the robustness of the optimal schedule in the presence of drug-resistant states, showing that although resistance reduces efficacy, the optimized regimen remains the best-performing option across multiple resistance scenarios (Fig. 4). We also demonstrated the use of the digital twin model for the identification of individualized optimal treatment schedules using patient-specific treatment-response data (Fig. 6). It is important to note that our optimization focused on the treatment cycle itself. Maintenance dosing beyond the cycle, which for many targeted agents is administered until disease progression, was not explicitly optimized and could be incorporated in future extensions of the framework.

One of the main strengths of our digital twin model is its adaptability. For example, we evaluated how an alternative AKT inhibitor (capivasertib) influences treatment efficacy simply by substituting the compound-specific PK model within the model (Fig. 5). A similar analysis could be performed for alternative EZH2 inhibitors to tazemetostat, or for other drugs targeting the same pathways, by incorporating the corresponding pharmacokinetic models.

More broadly, the digital twin model developed in this paper can be extended to investigate additional chromatin regulators that may influence tumor progression and treatment response. Because the digital twin integrates a mechanistic model describing the interaction between chromatin dynamics and cancer cell-state transitions, new epigenetic modifiers can be readily incorporated by modifying the underlying regulatory network to reflect their biological mechanisms of action. This enables systematic exploration of how different chromatin regulators could affect tumor dynamics and therapeutic outcomes, providing a flexible tool to generate and test mechanistic hypotheses about epigenetic interventions in cancer.

In terms of future extensions of the digital twin model, it is important to note that in the current chromatin modification circuit each nucleosome is assumed to carry either a repressive or an activating chromatin mark, without the possibility of an unmodified state. This simplifying assumption preserves the autocatalytic feedback loops underlying histone modification maintenance [21] while reducing model dimensionality. However, more elaborate circuits including intermediate or unmodified chromatin states may be necessary in contexts where additional chromatin regulatory mechanisms play a key role in therapy response.

In conclusion, this work provides a flexible digital twin model that can be used to understand the mechanistic basis of epigenetic therapies, predicting treatment responses, and rationally optimizing drug schedules. In this study, we illustrated the capabilities of the digital twin model in the context of TNBC, but the underlying modeling strategy is broadly generalizable and can be adapted to study other cancers in which chromatin modifications play a critical role in tumor progression and therapeutic response. Indeed, epigenetic therapies are being actively explored across multiple tumor types. For example, DNA methyltransferase inhibitors such as the FDA-approved agents azacitidine and decitabine are widely used in hematologic malignancies, such as acute myeloid leukemia [54, 55]. Histone deacetylase inhibitors, including vorinostat, romidepsin, belinostat, and panobinostat, have also been approved for the treatment of several lymphoid cancers [56, 57]. Beyond hematologic malignancies, epigenetic therapies are also being actively investigated in a variety of solid tumors, including, in addition to breast cancer, colorectal, pancreatic, and sarcoma cancers [58–61]. These developments highlight the relevance of extending this modeling approach more broadly across diverse cancer contexts. Overall, this digital twin model has the potential to support the development of novel, more effective, and more durable therapeutic strategies across different cancer types.

## Supplementary information

### S1 File

Supporting information file that contains full model details and additional analyses, including: the complete set of ODEs for the epigenetic TNBC progression model and its extension with drug-resistant states; posterior means of all model parameters; the derivation of the PK models for tazemetostat, ipatasertib, and capivasertib; and supplementary deterministic analyses exploring the impact of resistance mechanisms on treatment efficacy.

## Acknowledgments

Simone Bruno is a Damon Runyon Quantitative Biology Fellow and this work was supported by the Damon Runyon Cancer Research Foundation (DRQ-[2525]).

## Methods

### Cell culture and experimental assays

#### Cell Culture

Cell lines were purchased directly from ATCC or authenticated using STR analysis (Labcorp). SUM149PT cells (RRID:CVCL 3422) were cultured in DMEM/F12 (Gibco, 11330-057) and supplemented with 10% FBS, and 1X concentration of penicillin–streptomycin–glutamine (Gibco, 10378016). Cell lines were routinely found to have no mycoplasma contamination using Lonza MycoAlert PLUS Mycoplasma Detection Kit (Lonza, LT07-710). SUM149PT cells are derived from a female human breast carcinoma and all cell culture experiments were done with this cell line.

#### IncuCyte live-cell imaging

The protocol for live-cell imaging is the following: live-cell imaging was completed using IncuCyte live-cell imaging. SUM149PT cells were pretreated for 5 days with EZH2i (tazemetostat, SelleckChem, S7128) or DMSO (vehicle). Cells were then seeded at 5,000 cells per well in 96-well plates (Fisher, 08-772-225) and allowed to incubate overnight. Cells were then treated with EZH2i or AKTi (ipatasertib, Selleckchem, S2808) at varying concentrations in media containing NucLight Rapid Red Reagent (Sartorius, 4717, 1:1000) to label nuclei and Cytotox Green (Sartorius, 4633, 1:4000) to label cytotoxic cells. Plates were placed inside the IncuCyte machine and images were acquired every 2 hours for 5 days. The percentage of cytotoxic cells was determined using the integrated IncuCyte software by quantifying the overlap of NucLight- and Cytotox-green-positive cells divided by the total number of NucLight-positive cells. Four images per well were taken and each condition was seeded in triplicate. All of the experiments were completed at least three times.

### Data analysis

All data manipulation and plotting were performed in R (v4.3.2) [62] using the package `tidyverse` (v2.0.0) [63]. Experimental measurements originally provided as Excel files were imported in R and converted to `.Rda` objects for analysis. Data preprocessing, visualization, and preparation of input lists for the Stan model were carried out entirely in R. Parameter inference and posterior analysis were performed as described below, and posterior analyses and plotting were performed using custom R scripts.

### Parameter estimation in the epigenetic TNBC progression model

Bayesian inference [64] used to estimate parameters in our TNBC progression model was performed using the Stan programming language (v2.32.2) [65] in the R package `rstan` (v2.32.6) [66]. We used Hamiltonian Monte Carlo with the No-U-Turn Sampler (NUTS) [67] to fit the system of ODEs in (S.1), solved with Stan’s stiff solver `integrate ode bdf`. Broad Normal priors were assigned to all parameters to allow flexibility in estimation [68]. Experimental measurements of the percentage of dying cells were assumed to be normally distributed around the corresponding model predictions, with the observation-noise standard deviation estimated jointly with the model parameters. The quantified IncuCyte live-cell imaging data of *in vitro* TNBC cultures (Fig. 2B) were then used to estimate posterior distributions of the ODE model parameters (SI-Fig. S.10). Initial conditions for the percentage of dying cells were set equal to the experimentally measured mean percentage of dying cells at treatment initiation. Because measurements distinguishing basal-like and luminal-like cells were not available, all remaining viable cells were assumed to be in the basal-like state, which represents the predominant phenotype of TNBC cells [17]. Drug inputs were modeled according to the experimental design: AKTi was applied at *t* ≥ 5 *×* 24 hours, while EZH2i was applied continuously. Furthermore, both drug concentrations were assumed to remain approximately constant throughout the experiment (240 hours, i.e., 5 days of pretreatment followed by 5 days of treatment). In Fig. 2C, we present the posterior mean trajectory, computed as the average of trajectories simulated across posterior parameter samples, together with the experimental data for comparison.

Model accuracy was quantified by computing the Pearson correlation [69] between posterior predictive mean trajectories and experimental replicate-mean trajectories, together with the root mean squared error (RMSE) normalized by inter-replicate experimental variability, i.e., the average standard deviation across experimental replicates.

### *In silico* clinical trials

We simulated *in silico* clinical trials by generating a cohort of 500 virtual patients. Pharmacokinetics (PK) of tazemetostat and ipatasertib (or capivasertib) were modeled using a one-compartment model parameterized from published pharmacokinetic data and a published three-compartment model, respectively (see Sections “Development of the digital twin model”, “Impact of alternative AKT inhibitors or epigenetic modifiers on treatment response”, and SI-Sections S.3,S.4, and S.6). Inter-patient variability in PK parameters was incorporated by sampling individual PK parameters according to the published coefficients of variation or inter-individual variability estimates for each drug ([38, 49, 50] for tazemetostat, [39] for ipatasertib, and [53] for capivasertib). Inter-patient variability in pharmacodynamic (PD) parameters was introduced by sampling, for each virtual patient, PD parameters from the posterior distributions inferred in the “Parameter estimation and validation using *in vitro* data” section (Fig. 3A).

Each simulation was run for 26 days, a duration recommended by our clinical collaborators as a reasonable timeframe to capture meaningful treatment response. The primary outcome was defined as the percentage of dying cells at the end of the simulation (day 26), which we used as a proxy for treatment efficacy. For each tested regimen, we obtained the distribution of outcomes across the virtual cohort, allowing for direct comparisons between schedules.

To assess the robustness of the treatment schedule ranking, we additionally evaluated the selected regimens using two alternative efficacy metrics and an extended evaluation period. Specifically, we computed cell survival AUC, defined as the time integral of the surviving cell fraction over the simulation period, and the cumulative death induction, defined as the time integral of the model-predicted death induction rate. We also evaluated treatment efficacy, i.e., percentage of dying cells, after two treatment cycles (day 47).

### Genetic algorithm for treatment schedule optimization

In order to optimize treatment schedules, we use a genetic algorithm (GA) [51], a stochastic population-based search method inspired by natural selection. In a GA, an initial population of candidate solutions (here, drug administration schedules encoded as state vectors) is generated at random within specified constraints. At each iteration (generation), the fitness of each candidate is evaluated according to a predefined objective function. In our case, the objective function was defined as the mean percentage of dying cells at the end of the 26-day treatment cycle across the simulated virtual cohort. Candidate schedules exceeding predefined cumulative exposure limits for either EZH2i or AKTi (derived from the reference treatment regimen, SI-Fig. S.1) were assigned a large penalty, thereby excluding excessively toxic schedules from the optimization.

The next generation of candidates is then formed by selecting the top-performing regimens (elitism), applying random perturbations to some candidates (mutation), and recombining parts of two parent schedules to form new offspring (crossover). This process is repeated over multiple generations, allowing the algorithm to progressively refine the population toward higher-performing regimens.

All simulations were implemented in R (v4.3.2), using the package `GA` to perform the genetic algorithm search. The GA parameters were set to a population size of 70, with 70 generations, crossover probability 0.8, and mutation probability 0.2. The algorithm was terminated if no improvement in the best fitness value was observed for 30 consecutive generations, and the best-performing regimen at convergence was selected as the predicted optimal schedule.

In Fig. 3B, we show all treatment schedules evaluated by the GA for the tazemetostat (EZH2i) + ipatasertib (AKTi) regimen, and in SI-Fig. S.6 we show all schedules evaluated for the tazemetostat (EZH2i) + capivasertib (AKTi) regimen.

### Deterministic analyses

We performed deterministic analyses to study our TNBC progression model including drug-resistant states (Fig. 4A and SI-Figs. S.4,S.5). The ODE system (SI-Eq. (S.18)) was simulated using the package `deSolve` in R. The parameter values used for the analysis are provided in SI-Section S.5.

### Individualized epigenetic therapy optimization

To demonstrate how the digital twin model can be used to identify individualized optimal therapies, we performed a proof-of-concept computational study using synthetic patient-specific *in vitro* treatment response data. These data were generated as follows. For each virtual patient, we randomly sampled parameter values from the posterior distributions previously estimated from the population-level calibration (SI-Fig. S.10). We then assigned these parameter values to our TNBC progression ODE model, simulated treatment responses for different combinations and concentrations of the AKTi ipatasertib and the EZH2i tazemetostat, and added random measurement noise to the simulated trajectories. For each treatment condition, three experimental replicates were generated.

For patient-specific parameter estimation, we first identified *δ*_2_, *K*_*A*_, *K*_*E*_, and *δ*_1_ as the four patient-specific model parameters governing the response to AKT and EZH2 inhibition. These four parameters were selected because they directly regulate the strength (*δ*_1_ and *δ*_2_) and sensitivity (*K*_*A*_ and *K*_*E*_) of the pharmacodynamic response to the two drugs and were therefore expected to exhibit the greatest patient-to-patient variability. In contrast, the remaining parameters describe intrinsic properties of the TNBC progression model that were assumed to be conserved across patients. We then estimated these four patient-specific parameters using the same Bayesian inference framework described above, using the posterior distributions obtained from the population-level calibration (SI-Fig. S.10) as prior distributions, while fixing all remaining model parameters to their population-level posterior mean values. The patient-specific parameter values used for subsequent analyses were taken as the means of the corresponding estimated posterior distributions (SI-Fig. S.11).

Finally, the optimal patient-specific treatment schedule was identified by applying the same genetic algorithm optimization framework used for the population-level analyses, replacing the population model with the corresponding personalized model.

Pharmacokinetic parameters were not re-estimated during this study. Instead, the PK models were simulated using the previously estimated population-level mean parameter values. This choice reflects the fact that patient-specific pharmacokinetic measurements may not be available during the early stages of treatment optimization, whereas population pharmacokinetic models are routinely used to predict drug exposure using established population parameter estimates.

