## Supplementary information: S1 file for "A mechanistic digital twin model for epigenetic therapy optimization in triple-negative breast cancer"

### S.1 ODEs of the epigenetic TNBC progression model

In this section, we derive the ODEs associated with the epigenetic TNBC progression model introduced in the paper. The model tracks the dynamics of three cellular states: dying cells, luminal-like cells, and basal-like cells. Let $Y_1(t)$ denote the percentage of dying cells, $Y_8(t)$ the percentage of luminal-like cells, and $B(t)$ the percentage of basal-like cells. Because these three populations represent the total cell population in our system, we have the conservation law $Y_1(t)+Y_8(t)+B(t) = 100$, which allows us to reduce the system dimension by expressing the percentage of basal-like cells as function of the others, that is $B(t) = 100 - Y_1(t) - Y_8(t)$.

Two key transcription factors, GATA3 and BMF, regulate transitions between luminal-like and basal-like cell states. Their protein concentrations are given by $Y_2(t)$ and $Y_6(t)$, respectively. Furthermore, let $Y_4(t)$ and $Y_5(t)$ denote the concentrations of AKT and EZH2, respectively, and let $I_A$ and $I_E$ denote the concentrations of their corresponding inhibitors.

Concerning the chromatin modification circuit, it was constructed based on previous models of chromatin modification dynamics [1–3] developed by using the detailed characterizations of these modifications done over the past years [4–8]. For each gene (GATA3 and BMF), the basic modifiable unit is a nucleosome, and the total number of nucleosomes per gene is denoted by $\mathrm{D_{tot}}$. Each nucleosome can be modified either with a repressive (H3K27me3) or an activating (H3K4me3) histone modification. To reduce model complexity, we consider a simplified representation in which each nucleosome is either in an activating or repressive state, with no unmodified state. Under this assumption, the erasure of one modification coincides with the establishment of the opposing one, so that activation and repression dynamics are described through a single effective process. This simplification preserves the key mechanisms of chromatin regulation, including establishment, erasure, and positive feedback loops underlying maintenance, while simplifying the model.

Furthermore, we assume that $\mathrm{D_{tot}}$ is sufficiently large so that the number of nucleosomes with activating or repressive modifications can be treated as real-valued. In particular, we define $Y_3(t)$ as the number of nucleosomes with activating modifications for GATA3 and $Y_7(t)$ as the number of nucleosomes with activating modifications for BMF. The corresponding numbers of nucleosomes having repressive modifications are $R_G(t) = \mathrm{D_{tot}} - Y_3(t)$ and $R_B(t) = \mathrm{D_{tot}} - Y_7(t)$, based on the conservation of the total number of nucleosomes per gene. For simplicity, we assume that $\mathrm{D_{tot}}$ is the same for both genes. While this reduces the computational cost of our analyses, it does not affect the overall conclusions of our study. Since there is approximately one nucleosome per 200 bp ([9], Chapter 4) and an average gene spans 10,000–20,000 bp [10], $\mathrm{D_{tot}}$ can be considered on average between 50 and 100. In particular, in our analyses we set $\mathrm{D_{tot}} = 50$.

Concerning the parameters of our ODE model, we define them as follows:

- $\alpha_*$ terms, with $* = G, B, A, E$ ($\mu$M/h or $\mu$M/(h $\cdot$ nucleosome)) denote gene expression rates.
- $\delta_d$ ($\mathrm{h}^{-1}$) denotes the clearance rate of dying cells.
- $\delta_A^0$ and $\delta_E^0$ ($\mathrm{h}^{-1}$) denote the basal decay rates of AKT and EZH2.
- $\delta_A$ and $\delta_E$ ($\mathrm{h}^{-1}$) denote the drug-induced degradation rate of AKT and EZH2 through their respective inhibitors.
- $\delta_G$ and $\delta_B$ ($\mathrm{h}^{-1}$) denote the basal decay rates of GATA3 and BMF.
- $\delta_1$ ($\mathrm{h}^{-1}$) denotes the rate at which basal-like cells transition into the luminal-like state.

- $\delta_2$ ($\mathrm{h}^{-1}$) denotes the treatment-induced death rate of luminal-like cells. Furthermore, we multiply $\delta_2$ by an exponentially decaying factor $e^{-mt}$ to model loss of treatment effectiveness over time, with $m > 0$ ($\mathrm{h}^{-1}$) determining the rate at which this decline occurs.

- $\beta_G$ and $\beta_B$ ($\mathrm{h}^{-1}$) denote the basal rates of chromatin activation at the GATA3 and BMF genes.

- $\bar{\beta}_G$ and $\bar{\beta}_B$ ($1/(\mathrm{h}\cdot\mathrm{nucleosome})$) denote the rates of the read-write mechanism through which activating modifications enhance their own establishment for GATA3 and BMF.

- $\varepsilon_G$ and $\varepsilon_B$ ($\mathrm{h}^{-1}$) denote the basal rates of chromatin repression at the GATA3 and BMF genes.

- $\bar{\varepsilon}_G$ and $\bar{\varepsilon}_B$ ($\mathrm{h}^{-1}$) denote the rates of EZH2-mediated establishment of repressive mark H3K27me3 for GATA3 and BMF.

- $\hat{\varepsilon}_G$ and $\hat{\varepsilon}_B$ (1/nucleosome) are parameters encapsulating the strength of the autocatalytic process through which existing H3K27me3 marks enhance their own establishment by recruiting additional EZH2 for GATA3 and BMF.

The ODEs governing the epigenetic TNBC progression model can then be written as follows:

$$
\begin{aligned}
\frac{dY_1}{dt} &= \delta_2 e^{-mt}\frac{Y_6}{K_1+Y_6}Y_8 - \delta_d Y_1\left(1-\frac{Y_1}{100}\right)\\
\frac{dY_2}{dt} &= \alpha_G\frac{K_2}{K_2+Y_4}Y_3 - \delta_G Y_2\\
\frac{dY_3}{dt} &= (\beta_G+\bar{\beta}_G Y_3)(\mathrm{D_{tot}}-Y_3) - \left(\varepsilon_G+\bar{\varepsilon}_G\frac{Y_5}{K_3+Y_5}(1+\hat{\varepsilon}_G(\mathrm{D_{tot}}-Y_3))\right)Y_3\\
\frac{dY_4}{dt} &= \alpha_A - \delta_A\frac{I_A}{K_A+I_A}Y_4 - \delta_A^0 Y_4\\
\frac{dY_5}{dt} &= \alpha_E - \delta_E\frac{I_E}{K_E+I_E}Y_5 - \delta_E^0 Y_5\\
\frac{dY_6}{dt} &= \alpha_B\frac{K_6}{K_6+Y_4}Y_7 - \delta_B Y_6\\
\frac{dY_7}{dt} &= (\beta_B+\bar{\beta}_B Y_7)(\mathrm{D_{tot}}-Y_7) - \left(\varepsilon_B+\bar{\varepsilon}_B\frac{Y_5}{K_7+Y_5}(1+\hat{\varepsilon}_B(\mathrm{D_{tot}}-Y_7))\right)Y_7\\
\frac{dY_8}{dt} &= \delta_1\frac{Y_2}{K_5+Y_2}(100-Y_1-Y_8) - \left(\delta_2 e^{-mt}\frac{Y_6}{K_1+Y_6} - \delta_d\frac{Y_1}{100}\right)Y_8.
\end{aligned}
\tag{S.1}
$$

**Derivation of the epigenetic TNBC progression model** (S.1)
Let us start by deriving the ODEs for the cellular-state variables expressed as percentages. Let $\bar{B}(t)$, $\bar{Y}_8(t)$, and $\bar{Y}_1(t)$ denote the absolute numbers of basal-like, luminal-like, and dying cells, respectively. The cellular-state transition model, shown in Fig. 1B, can be summarized as

$$\bar{B} \longrightarrow \bar{Y}_8 \longrightarrow \bar{Y}_1 \longrightarrow \text{clearance},$$

in which the basal-to-luminal transition rate is $r_1(t) = \delta_1\frac{Y_2}{K_5+Y_2}$, the luminal-to-dying transition rate is $r_2(t) = \delta_2 e^{-mt}\frac{Y_6}{K_1+Y_6}$, and the dying-cell clearance rate is $\delta_d$. The corresponding ODEs can be written as

$$\frac{d\bar{B}}{dt} = -r_1(t)\bar{B}, \quad \frac{d\bar{Y}_8}{dt} = r_1(t)\bar{B} - r_2(t)\bar{Y}_8, \quad \frac{d\bar{Y}_1}{dt} = r_2(t)\bar{Y}_8 - \delta_d\bar{Y}_1. \tag{S.2}$$

Let us now define the total number of cells in the system as $N(t) = \bar{B}(t) + \bar{Y}_8(t) + \bar{Y}_1(t)$, whose dynamics are governed by the following ODE:

$$\frac{dN}{dt} = \frac{d\bar{B}}{dt} + \frac{d\bar{Y}_8}{dt} + \frac{d\bar{Y}_1}{dt} = -\delta_d\bar{Y}_1. \tag{S.3}$$

The percentages of $\bar{B}(t)$, $\bar{Y}_8$, and $\bar{Y}_1$ can then be defined as

$$B(t) = 100\frac{\bar{B}(t)}{N(t)}, \quad Y_8(t) = 100\frac{\bar{Y}_8(t)}{N(t)}, \quad Y_1(t) = 100\frac{\bar{Y}_1(t)}{N(t)}. \tag{S.4}$$

Furthermore, by definition we have $B(t) + Y_8(t) + Y_1(t) = 100$, which implies $B(t) = 100 - Y_1(t) - Y_8(t)$. Now, applying the quotient rule for derivatives, we obtain

$$\frac{dY_1}{dt} = 100\frac{d}{dt}\left(\frac{\bar{Y}_1}{N}\right) = 100\left(\frac{1}{N}\frac{d\bar{Y}_1}{dt} - \frac{\bar{Y}_1}{N^2}\frac{dN}{dt}\right). \tag{S.5}$$

Substituting in SI-Eq. (S.5) the expressions for $\frac{d\bar{Y}_1}{dt}$ (SI-Eq. (S.2)) and $\frac{dN}{dt}$ (SI-Eq. (S.3)) and the definitions of $Y_1$ and $Y_8$ (SI-Eq. (S.4)), we obtain

$$\begin{aligned}\frac{dY_1}{dt} &= 100\left(\frac{r_2(t)\bar{Y}_8 - \delta_d\bar{Y}_1}{N} + \frac{\delta_d\bar{Y}_1^2}{N^2}\right) = r_2(t)Y_8 - \delta_d Y_1 + \delta_d\frac{Y_1^2}{100} \\ &= r_2(t)Y_8 - \delta_d Y_1\left(1 - \frac{Y_1}{100}\right).\end{aligned} \tag{S.6}$$

Similarly, for the percentage of the luminal-like cells, $Y_8(t)$, we obtain

$$\frac{dY_8}{dt} = 100\frac{d}{dt}\left(\frac{\bar{Y}_8}{N}\right) = 100\left(\frac{1}{N}\frac{d\bar{Y}_8}{dt} - \frac{\bar{Y}_8}{N^2}\frac{dN}{dt}\right). \tag{S.7}$$

Substituting in SI-Eq. (S.7) the expressions for $\frac{d\bar{Y}_8}{dt}$ (SI-Eq. (S.2)) and $\frac{dN}{dt}$ (SI-Eq. (S.3)) and the definitions of $Y_1$, $Y_8$ and $B(t)$ (SI-Eq. (S.4)), we obtain

$$\begin{aligned}\frac{dY_8}{dt} &= 100\left(\frac{r_1(t)\bar{B} - r_2(t)\bar{Y}_8}{N} + \frac{\delta_d\bar{Y}_1\bar{Y}_8}{N^2}\right) = r_1(t)B - r_2(t)Y_8 + \delta_d\frac{Y_1Y_8}{100} \\ &= r_1(t)B - r_2(t)Y_8 + \delta_d\frac{Y_1Y_8}{100} = r_1(t)(100 - Y_1 - Y_8) - \left(r_2(t) - \delta_d\frac{Y_1}{100}\right)Y_8.\end{aligned} \tag{S.8}$$

Finally, substituting the expressions for $r_1(t)$ and $r_2(t)$, in SI-Eqs. (S.6) and (S.8), we obtain

$$\begin{aligned}\frac{dY_1}{dt} &= \delta_2 e^{-mt}\frac{Y_6}{K_1 + Y_6}Y_8 - \delta_d Y_1\left(1 - \frac{Y_1}{100}\right), \\ \frac{dY_8}{dt} &= \delta_1\frac{Y_2}{K_5 + Y_2}(100 - Y_1 - Y_8) - \left(\delta_2 e^{-mt}\frac{Y_6}{K_1 + Y_6} - \delta_d\frac{Y_1}{100}\right)Y_8.\end{aligned}$$

which correspond to the equations in the full ODE model (S.1).

We next derive the ODEs governing the molecular species and chromatin-state variables. In our model, $Y_4(t)$ and $Y_5(t)$ denote the concentrations of AKT and EZH2, respectively. Their dynamics are modeled through basal production, basal degradation, and inhibitor-induced degradation. For AKT, we assume a constant production rate $\alpha_A$, a basal degradation rate $\delta_A^0$, and an inhibitor-induced degradation rate that saturates with the concentration of the AKT inhibitor $I_A$, which we model through a Hill function [11]. The resulting ODE can be written as

$$\frac{dY_4}{dt} = \alpha_A - \delta_A \frac{I_A}{K_A + I_A} Y_4 - \delta_A^0 Y_4.$$

The dynamics of EZH2 are modeled similarly, yielding

$$\frac{dY_5}{dt} = \alpha_E - \delta_E \frac{I_E}{K_E + I_E} Y_5 - \delta_E^0 Y_5.$$

Concerning GATA3 and BMF, they are represented by $Y_2(t)$ and $Y_6(t)$, respectively. Furthermore, we assume that transcription occurs only from nucleosomes carrying activating histone modifications [1]. As a consequence, the production rate of GATA3 is proportional to $Y_3(t)$, i.e., the number of nucleosomes carrying activating modifications associated with the GATA3 gene. Finally, modeling the inhibitory effect of AKT on GATA3 expression through a decreasing Hill function [11], we obtain

$$\frac{dY_2}{dt} = \left(\alpha_G \frac{K_2}{K_2 + Y_4}\right) Y_3 - \delta_G Y_2.$$

We then analogously modeled the dynamics of BMF, obtaining

$$\frac{dY_6}{dt} = \left(\alpha_B \frac{K_6}{K_6 + Y_4}\right) Y_7 - \delta_B Y_6.$$

Let us now derive the ODEs associated with chromatin modification dynamics. For GATA3, since the number of nucleosomes associated with a gene is conserved and since we denoted this number by $\mathrm{D_{tot}}$, the quantity $\mathrm{D_{tot}} - Y_3(t)$ represents the number of nucleosomes carrying repressive modifications.

Activation of repressed nucleosomes is assumed to occur through two mechanisms: a basal activation process and a read-write positive-feedback mechanism mediated by existing activating modifications [1]. Conversely, activating modifications can be converted into repressive modifications through both basal repression and EZH2-mediated repression. The EZH2-dependent component is modeled through an increasing Hill function. Furthermore, we assume that existing repressive modifications can also enhance their own establishment through a read-write positive-feedback mechanism. Combining these effects, the effective repression rate is

$$\left(\varepsilon_G + \bar{\varepsilon}_G \frac{Y_5}{K_3 + Y_5}(1 + \hat{\varepsilon}_G(\mathrm{D_{tot}} - Y_3))\right) \tag{S.9}$$

Then, putting together the activation and repression terms, the ODE for $Y_3(t)$ can be written as

$$\frac{dY_3}{dt} = (\beta_G + \bar{\beta}_G Y_3)(\mathrm{D_{tot}} - Y_3) - \left(\varepsilon_G + \bar{\varepsilon}_G \frac{Y_5}{K_3 + Y_5}(1 + \hat{\varepsilon}_G(\mathrm{D_{tot}} - Y_3))\right) Y_3$$

Finally, the chromatin dynamics of BMF are modeled analogously. More precisely, denoting the number of nucleosomes carrying repressive modifications for BMF by $\mathrm{D_{tot}} - Y_7(t)$, the ODE for $Y_7(t)$ can be written as

$$\frac{dY_7}{dt} = (\beta_B + \bar{\beta}_B Y_7)(\mathrm{D_{tot}} - Y_7) - \left(\varepsilon_B + \bar{\varepsilon}_B \frac{Y_5}{K_7 + Y_5}(1 + \hat{\varepsilon}_B(\mathrm{D_{tot}} - Y_7))\right) Y_7.$$

### S.2 Parameter estimation: posterior means

Starting from the posterior distributions of the parameters (SI-Fig. S.10) of the epigenetic TNBC progression model (SI-Eq. (S.1)), obtained as described in the "Parameter estimation and validation using *in vitro* data" section and Methods, we computed the posterior mean of each parameter:

$$
\begin{aligned}
&\delta_2 = 3.799507 \times 10^{-1}\ \mathrm{h}^{-1}, \quad K_1 = 8.717916\ \mu\mathrm{M}, \quad \alpha_G = 1.173375\ \frac{\mu\mathrm{M}}{\mathrm{h}\cdot\mathrm{nucleosome}},\\
&K_2 = 1.203015 \times 10^{1}\ \mu\mathrm{M}, \quad \delta_G = 4.520690 \times 10^{-2}\ \mathrm{h}^{-1}, \quad \beta_G = 1.272923\ \mathrm{h}^{-1},\\
&\bar{\varepsilon}_G = 1.108645 \times 10^{-1}\ \mathrm{h}^{-1}, \quad K_3 = 9.394812\ \mu\mathrm{M}, \quad \alpha_A = 1.566855 \times 10^{-4}\ \frac{\mu\mathrm{M}}{\mathrm{h}},\\
&\delta_A = 4.560950 \times 10^{-1}\ \mathrm{h}^{-1}, \quad K_A = 1.952322 \times 10^{1}\ \mu\mathrm{M}, \quad \alpha_E = 1.333967\ \frac{\mu\mathrm{M}}{\mathrm{h}},\\
&\delta_E = 1.256836 \times 10^{-1}\ \mathrm{h}^{-1}, \quad K_E = 3.627773 \times 10^{-1}\ \mu\mathrm{M}, \quad \delta_d = 4.110200 \times 10^{-3}\ \mathrm{h}^{-1}, \qquad \text{(S.10)}\\
&m = 1.227741 \times 10^{-2}\ \mathrm{h}^{-1}, \quad \delta_1 = 1.603587 \times 10^{-1}\ \mathrm{h}^{-1}, \quad K_5 = 1.003410 \times 10^{1}\ \mu\mathrm{M},\\
&\alpha_B = 1.619134\ \frac{\mu\mathrm{M}}{\mathrm{h}\cdot\mathrm{nucleosome}}, \quad K_6 = 1.640542 \times 10^{-5}\ \mu\mathrm{M}, \quad \delta_B = 9.486298 \times 10^{-2}\ \mathrm{h}^{-1},\\
&\beta_B = 4.403414 \times 10^{-1}\ \mathrm{h}^{-1}, \quad \bar{\varepsilon}_B = 2.477224 \times 10^{-1}\ \mathrm{h}^{-1}, \quad K_7 = 1.341020 \times 10^{1}\ \mu\mathrm{M},\\
&\bar{\beta}_G = 1.360458\ \frac{1}{\mathrm{h}\cdot\mathrm{nucleosome}}, \quad \hat{\varepsilon}_G = 1.336714\frac{1}{\mathrm{nucleosome}}, \quad \bar{\beta}_B = 1.655805 \times 10^{-1}\ \frac{1}{\mathrm{h}\cdot\mathrm{nucleosome}},\\
&\hat{\varepsilon}_B = 2.233409\frac{1}{\mathrm{nucleosome}}, \quad \varepsilon_G = 1.002968\ \mathrm{h}^{-1}, \quad \varepsilon_B = 1.147803 \times 10^{-1}\ \mathrm{h}^{-1},\\
&\delta_A^0 = 4.520690 \times 10^{-2}\ \mathrm{h}^{-1}, \quad \delta_E^0 = 4.520690 \times 10^{-2}\ \mathrm{h}^{-1}.
\end{aligned}
$$

### S.3 PK model of tazemetostat

To model the pharmacokinetics (PK) of tazemetostat, we used the clinical pharmacology information reported in the FDA prescribing information together with published clinical pharmacokinetic analyses of tazemetostat [12–14], which provide estimates of oral bioavailability, absorption kinetics, apparent clearance, apparent volume of distribution, and evidence of time-dependent autoinduction. Although these sources report several summary pharmacokinetic quantities, they do not provide sufficiently detailed concentration-time data or estimates of inter-compartmental transfer rates required to identify and parameterize a multi-compartment model. Therefore, introducing additional compartments would require assigning values to parameters that cannot be directly obtained from the available clinical data. For this reason, we adopted a one-compartment model with first-order absorption and elimination, representing the simplest model structure supported by the available pharmacokinetic information.

In order to introduce the ODE model, let us denote the amount of drug in the gastrointestinal compartment as $A_{\mathrm{GI}}(t)$ and the plasma concentration of tazemetostat as $C_p(t)$. The PK model can be written as follows:

$$\frac{dA_{\mathrm{GI}}}{dt} = -k_a A_{\mathrm{GI}}, \quad \frac{dC_p}{dt} = \frac{k_a}{V_d} A_{\mathrm{GI}} - k_e(t) C_p, \qquad \text{(S.11)}$$

in which $k_a$ denotes the first-order absorption rate constant, $k_e(t)$ denotes the time-dependent elimination rate constant, and $V_d$ denotes the apparent volume of distribution.

To account for the reported absolute oral bioavailability of approximately (33%), each administered dose $D$ was multiplied by a factor $F = 0.33$ before entering the gastrointestinal compartment. Thus, immediately following a dosing event, the amount in the gastrointestinal compartment was increased according to $A_{\mathrm{GI}}(t^+) = A_{\mathrm{GI}}(t^-) + F \cdot D$.

The available clinical pharmacokinetic studies indicate that tazemetostat undergoes time-dependent autoinduction of metabolism, resulting in an increase in apparent oral clearance from approximately $CL_0/F = 126$ L/h after the first dose to $CL_{ss}/F = 274$ L/h at steady state, which is reached after approximately 15 days of treatment [12, 13]. To capture this behavior within our one-compartment model, we modeled the apparent clearance as

$$CL(t) = CL_0 + (CL_{ss} - CL_0)\left(1 - e^{-k_{\mathrm{ind}} t}\right), \tag{S.12}$$

where $CL_0$ and $CL_{ss}$ denote the apparent oral clearances immediately after treatment initiation and at steady state, respectively, and $k_{\mathrm{ind}}$ represents the autoinduction rate constant. Since steady state is reached after approximately 15 days [12–14], we selected

$$k_{\mathrm{ind}} = \frac{3}{15 \times 24} \approx 8.3 \times 10^{-3}\ \mathrm{h}^{-1}, \tag{S.13}$$

such that approximately 95% of the increase in clearance is achieved after 15 days. The corresponding time-dependent elimination rate constant was therefore computed as

$$k_e(t) = \frac{CL(t)}{V_d}, \tag{S.14}$$

where the apparent volume of distribution was fixed at $V_d/F = 681$ [14], as it yielded simulated peak plasma concentrations consistent with the reported clinical pharmacokinetic data [12–14].

Finally, the first-order absorption rate constant was fixed at $k_a = 2.42\ \mathrm{h}^{-1}$ [14], which is consistent with the reported median time to reach the peak plasma concentration ($T_{\max} = 1 - 2$ h) [12, 13].

To account for inter-individual pharmacokinetic variability, patient-specific values of the apparent volume of distribution ($V_d$), the apparent oral clearance following the first dose ($CL_0$), and the steady-state apparent oral clearance ($CL_{ss}$) were sampled from log-normal distributions [15]. Specifically, each parameter $P_i$ was generated according to

$$P_i = P\, e^{\eta_i}, \qquad \eta_i \sim \mathcal{N}(0, \omega_P^2), \tag{S.15}$$

where $P$ denotes the nominal value of the corresponding parameter and $\omega_P$ was computed from the reported coefficient of variation (CV) per parameter, $\mathrm{CV}_P$, as

$$\omega_P = \sqrt{\ln\left(1 + \mathrm{CV}_P^2\right)}, \tag{S.16}$$

with $\mathrm{CV}_P$ expressed as a fraction. The reported CVs of 46% for $V_d$ and 49% for clearance were taken from the FDA prescribing information [13]. Since separate estimates of inter-individual variability for $CL_0$ and $CL_{ss}$ were not available, both clearance parameters were assumed to share the reported clearance CV. Furthermore, for each virtual patient, the first-order absorption rate constant ($k_a$) and autoinduction rate constant ($k_{\mathrm{ind}}$) were fixed at their population values, while the time-dependent elimination rate constant was computed according to

$$k_e(t) = \frac{CL(t)}{V_d}, \tag{S.17}$$

using the sampled values of $V_d$, $CL_0$, and $CL_{ss}$. To ensure physiologically consistent autoinduction, the sampled steady-state clearance was constrained to satisfy $CL_{ss} \geq CL_0$ for every virtual patient, reflecting the clinically observed increase in clearance over time [14]. This procedure generated a heterogeneous virtual population while ensuring consistency with the reported clinical PK quantities.

For each virtual patient, the resulting PK model was simulated to obtain the corresponding plasma concentration profile of tazemetostat over time.

### S.4 PK model of ipatasertib

To model the pharmacokinetics (PK) of ipatasertib, we used the population PK model reported in [16]. Based on the analysis of clinical PK data, the authors identified a three-compartment model with first-order elimination and sequential zero- and first-order absorption as the model that best described the observed ipatasertib pharmacokinetics.

In order to introduce the ODE model, let us denote the amount of drug in the absorption depot compartment by $A_D(t)$, the amount of drug in the central compartment by $A_C(t)$, and the amounts of drug in the first and second peripheral compartments by $A_{P1}(t)$ and $A_{P2}(t)$, respectively. The PK model can then be written as

$$\frac{dA_D}{dt} = R_{\text{in}}(t) - k_a A_D, \quad \frac{dA_C}{dt} = k_a A_D - \frac{CL}{V_2} A_C - \frac{Q_3}{V_2} A_C + \frac{Q_3}{V_3} A_{P1} - \frac{Q_4}{V_2} A_C + \frac{Q_4}{V_4} A_{P2},$$
$$\frac{dA_{P1}}{dt} = \frac{Q_3}{V_2} A_C - \frac{Q_3}{V_3} A_{P1}, \quad \frac{dA_{P2}}{dt} = \frac{Q_4}{V_2} A_C - \frac{Q_4}{V_4} A_{P2},$$

in which $CL$ denotes the clearance, $V_2$, $V_3$, and $V_4$ denote the central and peripheral volumes of distribution, respectively, $Q_3$ and $Q_4$ denote the inter-compartmental clearances, and $k_a$ denotes the first-order absorption rate constant.

Drug administration was represented using a zero-order input process. Denoting the administered dose by $D$ and the duration of zero-order absorption by $t_0$, the input rate was defined as

$$R_{\text{in}}(t) = \frac{F_I D}{t_0},$$

during the interval $[t_{\text{dose}}, , t_{\text{dose}} + t_0]$, and zero otherwise.

The parameter values were taken directly from [16], namely $CL = 162$ L/h, $V_2 = 1230$ L, $V_3 = 2590$ L, $V_4 = 4340$ L, $Q_3 = 76.6$ L/h, $Q_4 = 3.26$ L/h, $k_a = 1.84$ h$^{-1}$, $t_0 = 0.348$ h.

Finally, the concentration driving the downstream pharmacodynamic model was taken as the central-compartment concentration, $C_{\text{AKTi}}(t) = A_C(t)/V_2$.

To account for inter-individual PK variability, patient-specific values of $CL$, $k_a$, and $t_0$ were sampled from log-normal distributions [15] according to the inter-individual variability (IIV) reported in [16]. Specifically, each parameter $P_i$ was sampled according to (S.15), in which $P$ denotes the nominal value of the corresponding parameter, reported above [16], and $\omega^2_{CL} = 0.0729$, $\omega^2_{k_a} = 1.74$, and $\omega^2_{t_0} = 2.23$ [16]. Furthermore, the PK model reported in [16] also quantifies inter-individual variability in relative bioavailability. Specifically, since individual body weights were not available, the body-weight effect was neglected by setting $\text{Weight}_i/\text{Weight}_{\text{med}} = 1$, yielding the following expression for the patient-specific relative bioavailability:

$$F_{I,i}(t) = (1 + \theta_{FI,MD} MD(t))\, e^{\eta_{FI,i}},$$

in which, following [16], the multiple-dosing indicator was defined as $MD(t) = 0$ for $t < 48$ h and $MD(t) = 1$ for $t \geq 48$ h, $\theta_{FI,MD} = 0.201$ $\eta_{FI,i} \sim \mathcal{N}(0, \omega^2_{FI})$, with $\omega^2_{FI} = 0.139$.

Finally, the PK analysis reported in [16] also characterized the active metabolite M1 (G-037720). However, we did not model M1 explicitly in the present study. This is because Yoshida et al. [16] reported that M1 is pharmacologically active but approximately 2–4-fold less potent than ipatasertib and it exhibits substantially lower systemic exposure, with a steady-state metabolite-to-parent AUC ratio of approximately 0.398. Therefore, the relative contribution of M1 to overall AKT inhibition can be approximated as 0.398/(2-4) = 0.10-0.20, indicating that M1 is expected to account for only about 10–20% of the total pharmacological activity. Then, while incorporating a dedicated PK model for M1 would substantially increase model complexity, it would provide only a limited contribution to overall target inhibition. This is the reason why only the parent compound ipatasertib was considered in the current model.

### S.5 ODEs of the epigenetic TNBC progression model including drug-resistant states

Here, we extended our original TNBC model (SI-Eq. S.1) to include a drug-resistance mechanism. In particular, we assume that cells may enter a drug-resistant state either from the basal-like cell state or from the luminal-like cell state (Fig. 4A). Let $Y_9(t)$ denote the percentage of resistant cells originating from the basal-like population and $Y_{10}(t)$ the percentage originating from the luminal-like population. In this extended model, the total cell population satisfies $Y_1(t) + Y_8(t) + Y_9(t) + Y_{10}(t) + B(t) = 100$, which allows us to express the basal-like population as $B(t) = 100 - Y_1(t) - Y_8(t) - Y_9(t) - Y_{10}(t)$.

Furthermore, let us introduce the following new parameters, associated with the resistance mechanisms:

- $a_1$ ($a_2$) denotes the transition rate from basal-like (luminal-like) cells to resistant cells.
- $b_1$ ($b_2$) denotes the death rate of resistant cells originating from basal-like (luminal-like) cells. Furthermore, to ensure that resistant cells remain less susceptible to treatment-induced death than luminal-like cells at all times, we multiply $b_1$ ($b_2$) by the same time-dependent exponential decay factor $e^{-mt}$. Without this adjustment, the global decline in treatment efficacy would eventually make sensitive cells die more slowly than resistant cells, which is biologically implausible.

The ODEs governing the TNBC progression model including drug-resistant states can then be written as follows:

$$
\begin{aligned}
\frac{dY_1}{dt} &= \delta_2 e^{-mt} \frac{Y_6}{K_1 + Y_6} Y_8 - \delta_d Y_1 \left(1 - \frac{Y_1}{100}\right) + b_1 e^{-mt} Y_9 + b_2 e^{-mt} Y_{10} \\
\frac{dY_2}{dt} &= \alpha_G \frac{K_2}{K_2 + Y_4} Y_3 - \delta_G Y_2 \\
\frac{dY_3}{dt} &= (\beta_G + \bar{\beta}_G Y_3)(\mathrm{D_{tot}} - Y_3) - \left(\varepsilon_G + \bar{\varepsilon}_G \frac{Y_5}{K_3 + Y_5}(1 + \hat{\varepsilon}_G(\mathrm{D_{tot}} - Y_3))\right) Y_3 \\
\frac{dY_4}{dt} &= \alpha_A - \delta_A \frac{I_A}{K_A + I_A} Y_4 - \delta_A^0 Y_4 \\
\frac{dY_5}{dt} &= \alpha_E - \delta_E \frac{I_E}{K_E + I_E} Y_5 - \delta_E^0 Y_5 \\
\frac{dY_6}{dt} &= \alpha_B \frac{K_6}{K_6 + Y_4} Y_7 - \delta_B Y_6 \\
\frac{dY_7}{dt} &= (\beta_B + \bar{\beta}_B Y_7)(\mathrm{D_{tot}} - Y_7) - \left(\varepsilon_B + \bar{\varepsilon}_B \frac{Y_5}{K_7 + Y_5}(1 + \hat{\varepsilon}_B(\mathrm{D_{tot}} - Y_7))\right) Y_7 \\
\frac{dY_8}{dt} &= \delta_1 \frac{Y_2}{K_5 + Y_2}(100 - Y_1 - Y_8 - Y_9 - Y_{10}) - \left(a_2 + \delta_2 e^{-mt} \frac{Y_6}{K_1 + Y_6} - \delta_d \frac{Y_1}{100}\right) Y_8 \\
\frac{dY_9}{dt} &= a_1(100 - Y_1 - Y_8 - Y_9 - Y_{10}) - (b_1 e^{-mt} - \delta_d \frac{Y_1}{100}) Y_9 \\
\frac{dY_{10}}{dt} &= a_2 Y_8 - (b_2 e^{-mt} - \delta_d \frac{Y_1}{100}) Y_{10}.
\end{aligned}
\tag{S.18}
$$

In the deterministic analysis conducted, we set $a_1 = A$ h$^{-1}$ and $a_2 = p_1 A$ h$^{-1}$, where $p_1$ is a dimensionless parameter scaling the ratio between $a_2$ and $a_1$. Similarly, we set $b_1 = B$ h$^{-1}$ and $b_2 = p_2 B$ h$^{-1}$, where $p_2$ is a dimensionless parameter scaling the ratio between $b_2$ and $b_1$.

We first simulated the ODEs in (S.18) for different values of $A$ and $B$, while keeping $p_1 = p_2 = 1$ and fixing all remaining parameters equal to the means of the posterior distributions previously estimated (SI–Eq. (S.10)) (SI-Fig. S.4). It is possible to observe that, for high values of $B$, the treatment has an equal or even greater effect in the model including resistant states than in the original model. This occurs because, when $B$ increases, the death rate of resistant cells increases until it can become higher than the death rate of sensitive cells. Although this scenario is mathematically possible, it is biologically implausible. If $A$ is low, then the treatment has essentially the same efficacy in the model with resistance as in the model without resistance. This is because, as $A$ decreases, fewer cells transition to the resistant state, resulting in a smaller fraction of resistant cells and therefore a smaller impact on treatment efficacy.

Thus, in the two-parameter regimes described above, the impact of resistance is low. In contrast, when $B$ is low (small resistant-cell death rate) and $A$ is high (many cells transition to a resistant state), the impact of resistance on treatment efficacy becomes substantial (high resistance) (SI-Fig. S.4). Overall, this analysis indicates that, within our modeling framework, the system corresponds to a case of "no resistance" when there are no transitions into resistant states, i.e., $a_1 = a_2 = b_1 = b_2 = 0$. It corresponds to a case of "high resistance" when resistant cells accumulate due to sufficiently large transition rates into resistance and sufficiently small death rates of resistant cells. To provide a quantitative operational definition, we define the high-resistance regime as the parameter region in which the percentage of dying cells decreases by at least 30% relative to the no-resistance case. For the posterior mean parameter values previously estimated for our epigenetic TNBC progression model, this regime corresponds to $a_1, a_2 \geq 0.01$ h$^{-1}$ and $b_1, b_2 \leq 0.01$ h$^{-1}$, excluding the boundary case in which all four parameters are simultaneously equal to 0.01 h$^{-1}$. All remaining biologically plausible parameter combinations correspond to a "low resistance" regime. It is important to note that these numerical thresholds are model-dependent and arise from our specific formulation of resistance dynamics and the posterior parameter values estimated in this study.

We then investigated the effects of $p_1$ and $p_2$ on treatment efficacy (SI-Fig. S.5). Our results show that differences in the transition rates to the resistant state from different cell states have a greater impact on treatment efficacy than differences in the death rates of the corresponding resistant states.

Finally, for the simulations showed in Fig. 4B, we consider $a_1 = a_2 = b_1 = b_2 = 0$ (no resistance), $a_1 = a_2 = 0.001$ h$^{-1}$; $b_1 = b_2 = 0.005$ h$^{-1}$ (low resistance), and $a_1 = a_2 = 0.01$ h$^{-1}$; $b_1 = b_2 = 0.005$ h$^{-1}$ (high resistance). All the other parameters, we set their values to the means of the posterior distributions previously estimated (SI-Eq. (S.10)).

### S.6 PK model of capivasertib

To model the pharmacokinetics (PK) of capivasertib, we used the population PK model reported in [17]. Based on the analysis of clinical PK data, Fernandez-Teruel et al. identified a three-compartment model with parallel first-order and zero-order absorption and time-dependent, dose-dependent apparent clearance. Since the absolute oral bioavailability was not available, all apparent parameters reported as $CL/F$, $V/F$, and $Q/F$ were used directly in the model.

In order to introduce the ODE model, let us denote the amounts of drug in the zero-order and first-order absorption pathways by $A_{D,0}(t)$ and $A_{D,1}(t)$, respectively,the amount of drug in the central compartment by $A_C(t)$, and the amounts of drug in the two peripheral compartments by $A_{P1}(t)$ and $A_{P2}(t)$, respectively. Denoting the administered dose by $D$, a fraction $0.2D$ was assigned to the zero-order absorption pathway and a fraction $0.8D$ to the first-order absorption pathway [17]. Furthermore, denoting the duration of zero-order absorption by $t_0$, the zero-order input rate was defined as

$$R_0(t) = \frac{0.2D}{t_0},$$

during the interval $[t_{\text{dose}},, t_{\text{dose}} + t_0]$, and zero otherwise. The PK model can then be written as

$$\begin{aligned}
\frac{dA_{D,0}}{dt} &= -R_0(t), \quad \frac{dA_{D,1}}{dt} = -k_a A_{D,1}, \\
\frac{dA_C}{dt} &= k_a A_{D,1} + R_0(t) - \frac{CL(t)}{V_2} A_C - \frac{Q_3}{V_2} A_C + \frac{Q_3}{V_3} A_{P1} - \frac{Q_4}{V_2} A_C + \frac{Q_4}{V_4} A_{P2}, \\
\frac{dA_{P1}}{dt} &= \frac{Q_3}{V_2} A_C - \frac{Q_3}{V_3} A_{P1}, \quad \frac{dA_{P2}}{dt} = \frac{Q_4}{V_2} A_C - \frac{Q_4}{V_4} A_{P2}.
\end{aligned}$$

The apparent clearance was modeled as time dependent according to [17]:

$$CL(t) = CL_0 \left[ 1 - \frac{e^{(I_{\max}(D))} t^5}{T_{50}^5 + t^5} \right],$$

in which $CL_0$ is the initial apparent clearance, $T_{50}$ is the time at which inhibition of clearance is half-maximal, and the dose-dependent maximal inhibition term, $I_{\max}(D)$, was defined as

$$I_{\max}(D) = I_{\max} \left[ 1 + (D - 480) I_{\max,\text{dose}} \right].$$

The parameter values were taken directly from [17], namely

$$\begin{aligned}
CL_0 = 62.2 \text{ L/h}, \ V_2 = 47.9 \text{ L}, \ V_3 = 113 \text{ L}, \ V_4 = 94.7 \text{ L}, \ Q_3 = 2.66 \text{ L/h}, \ Q_4 = 21.8 \text{ L/h}, \\
k_a = 0.417 \text{ h}^{-1}, \ t_0 = 45.1 \text{ h}, \ T_{50} = 67.4 \text{ h}, \ I_{\max} = -1.54, \ I_{\max,\text{dose}} = -0.00183 \text{ mg}^{-1}.
\end{aligned} \tag{S.19}$$

Finally, the concentration driving the downstream pharmacodynamic model was taken as the central-compartment concentration, $C_{\text{AKTi}}(t) = A_C(t)/V_2$.

To account for inter-individual PK variability, patient-specific values of $CL_0$, $V_2$, $I_{\max}$, $k_a$, and $t_0$ were sampled from log-normal distributions [15]. Specifically, each parameter $P_i$ was sampled according to (S.15), in which $P$ denotes the nominal value of the corresponding parameter, reported in (S.19), and $\omega_P$ was computed from the reported CV per parameter, $\text{CV}_P$, as (S.16), with $\text{CV}_P$ expressed as a fraction. The coefficients of variation were taken directly from [17], namely 39.3% for $CL_0$, 114% for $V_2$, 70.6% for $I_{\max}$, and 15% (fixed) for both $k_a$ and $t_0$.

### S.7 Supplementary figures

**Reference schedule**

| | Pretreat. (5 days) | Week 1 | Week 2 | Week 3 | Total |
|---|---|---|---|---|---|
| EZH2i | 800 mg BID | 800 mg BID | 800 mg BID | 800 mg BID | 41600 mg |
| AKTi | | 400 mg BID 4 days on 3 days off | 400 mg BID 4 days on 3 days off | 400 mg BID 4 days on 3 days off | 9600 mg |

**Candidate schedule – variable dose levels**

| | Pretreat. (5 days) | Week 1 | Week 2 | Week 3 | Total |
|---|---|---|---|---|---|
| EZH2i | $T_{pt}$ mg BID | $T_{w1}$ mg BID | $T_{w2}$ mg BID | $T_{w3}$ mg BID | < 41600 mg |
| AKTi | | $I_{w1}$ mg BID $W_1$ days on | $I_{w2}$ mg BID $W_2$ days on | $I_{w3}$ mg BID $W_3$ days on | < 9600 mg |

Figure S.1: **Treatment schedules evaluated by the genetic algorithm.** Summary of the reference schedule and the structure of candidate schedules, including all dosing constraints. The inhibitors considered are tazemetostat as EZH2i and ipatasertib or capivasertib as AKTi.

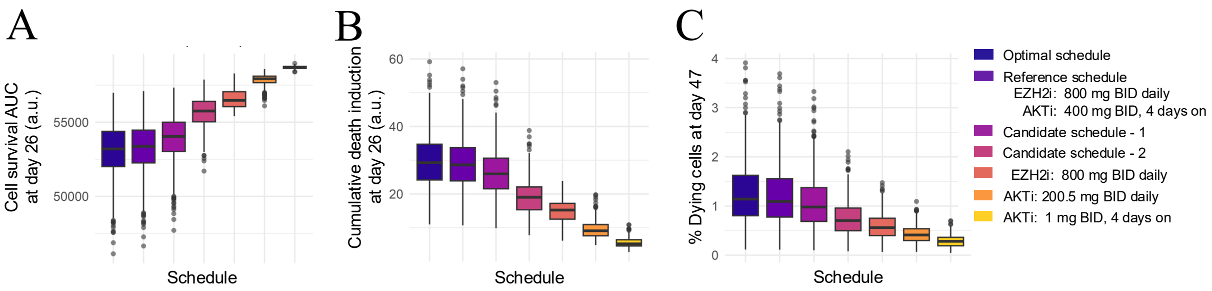


Figure S.2: **Additional evaluation metrics confirm the robustness of the optimized treatment schedule for the tazemetostat - ipatasertib combination. A** Box plots of regimen efficacy quantified using the cell survival area under the curve (AUC) at the end of one treatment cycle (day 26) (see Methods). **B** Box plots of regimen efficacy quantified using cumulative death induction at the end of one treatment cycle (day 26) (see Methods). **C** Box plots of regimen efficacy quantified using % of dying cells after two treatment cycles (day 47). For all panels, boxes represent the middle 50% of the data, with the median shown as a horizontal line. Whiskers extend to the minimum and maximum values excluding outliers, which are shown as dots. Each regimen is represented with a different color and its results are based on N = 500 simulations, corresponding to 500 *in silico* patients. The optimal schedule and candidate schedules 1 and 2 are the same as introduced in Fig. 3E.

A

**Discretized optimal schedule**

| | Pretreat. (5 days) | Week 1 | Week 2 | Week 3 | Total |
|---|---|---|---|---|---|
| Tazemetostat (EZH2i) | 600 mg BID | 600 mg BID | 600 mg BID | 400 mg BID | 28400 mg |
| Ipatasertib (AKTi) | | 400 mg BID 6 days on 1 days off | 300 mg BID 6 days on 1 days off | 200 mg BID 4 days on 3 days off | 10000 mg |

**Discretized candidate schedule – 1**

| | Pretreat. (5 days) | Week 1 | Week 2 | Week 3 | Total |
|---|---|---|---|---|---|
| Tazemetostat (EZH2i) | 400 mg BID | 800 mg BID | 600 mg BID | 400 mg BID | 29200 mg |
| Ipatasertib (AKTi) | | 300 mg BID 5 days on 2 days off | 300 mg BID 4 days on 3 days off | 100 mg BID 2 days on 5 days off | 5800 mg |

**Discretized candidate schedule – 2**

| | Pretreat. (5 days) | Week 1 | Week 2 | Week 3 | Total |
|---|---|---|---|---|---|
| Tazemetostat (EZH2i) | 400 mg BID | 400 mg BID | 400 mg BID | 0 mg BID | 15200 mg |
| Ipatasertib (AKTi) | | 100 mg BID 5 days on 2 days off | 300 mg BID 2 days on 5 days off | 200 mg BID 3 days on 4 days off | 3400 mg |

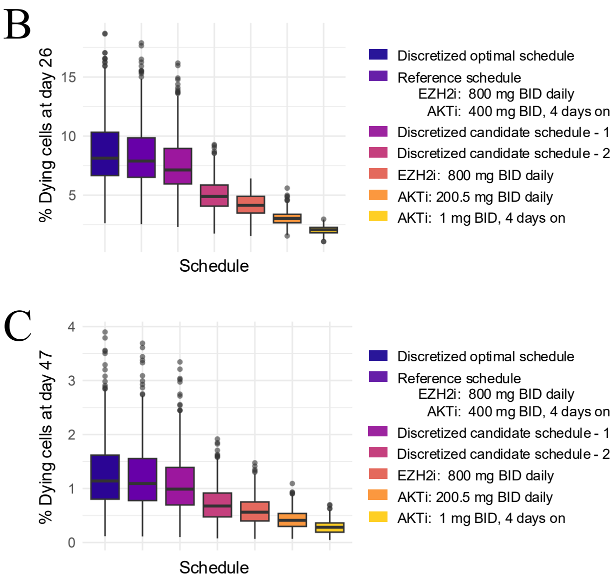


Figure S.3: **Evaluation of clinically feasible discretized treatment schedules for the tazemetostat - ipatasertib combination. A** Summary of the optimal schedule and candidate schedules 1 and 2, introduced in Fig. 3E, converted into clinically feasible regimens by rounding each dose to the nearest allowable dose level. Allowed doses are 0, 200, 400, 600, and 800 mg BID for tazemetostat and 0, 100, 200, 300, and 400 mg BID for ipatasertib. **B** Box plots of regimen efficacy quantified using % of dying cells after one treatment cycle (day 26) for the discretized schedules shown in panel A and the corresponding reference schedules. **C** Box plots of regimen efficacy quantified as % of dying cells after two treatment cycles (day 47) for the same regimens. For panels B and C, boxes represent the middle 50% of the data, with the median shown as a horizontal line. Whiskers extend to the minimum and maximum values excluding outliers, which are shown as dots. Each regimen is represented with a different color and its results are based on N = 500 simulations, corresponding to 500 *in silico* patients.

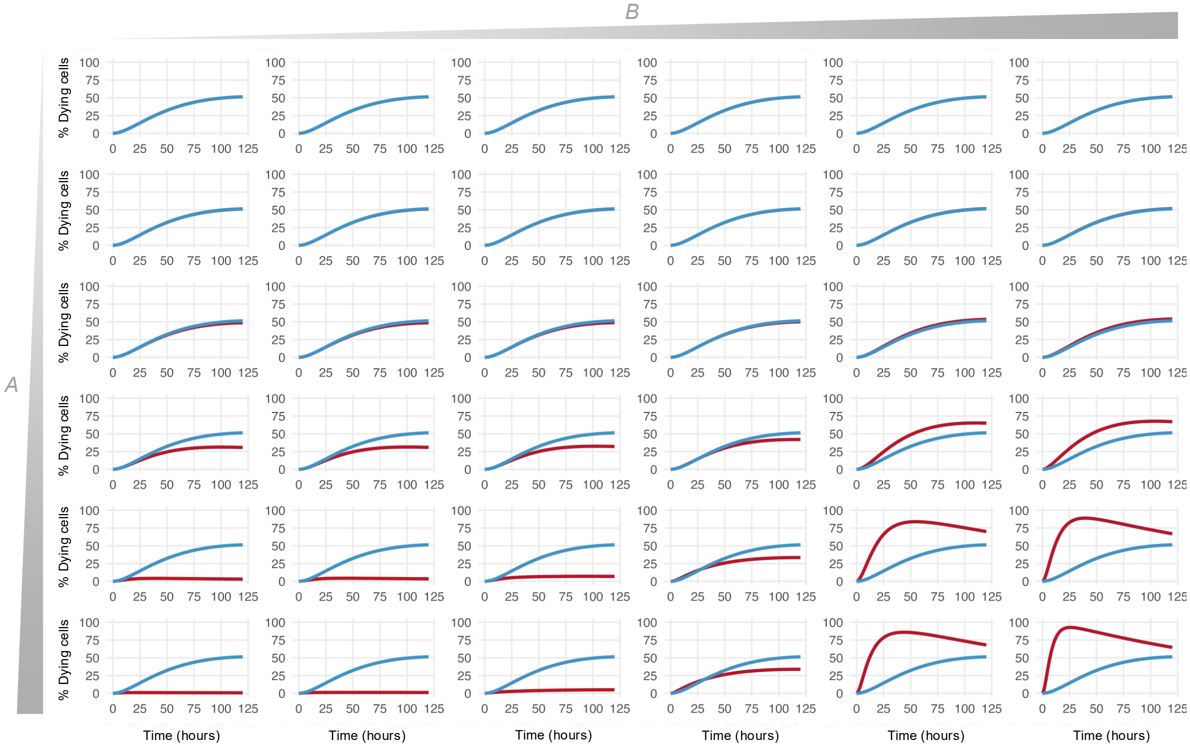


Figure S.4: **Effect of drug-resistant cell states on treatment efficacy.** Simulations of the TNBC ODE model including drug-resistant states (SI–Eq. S.18) for $A = 0, 0.0001, 0.001, 0.01, 0.1, 0.25$ $\mathrm{h}^{-1}$, $B = 0, 0.0001, 0.001, 0.01, 0.1, 0.25$ $\mathrm{h}^{-1}$, and $p_1 = p_2 = 1$ (red curves). The blue curve corresponds to the simulation of the TNBC model without drug-resistant states (i.e., $A = B = 0$). The complete list of parameter values is provided in SI–Section S.5.

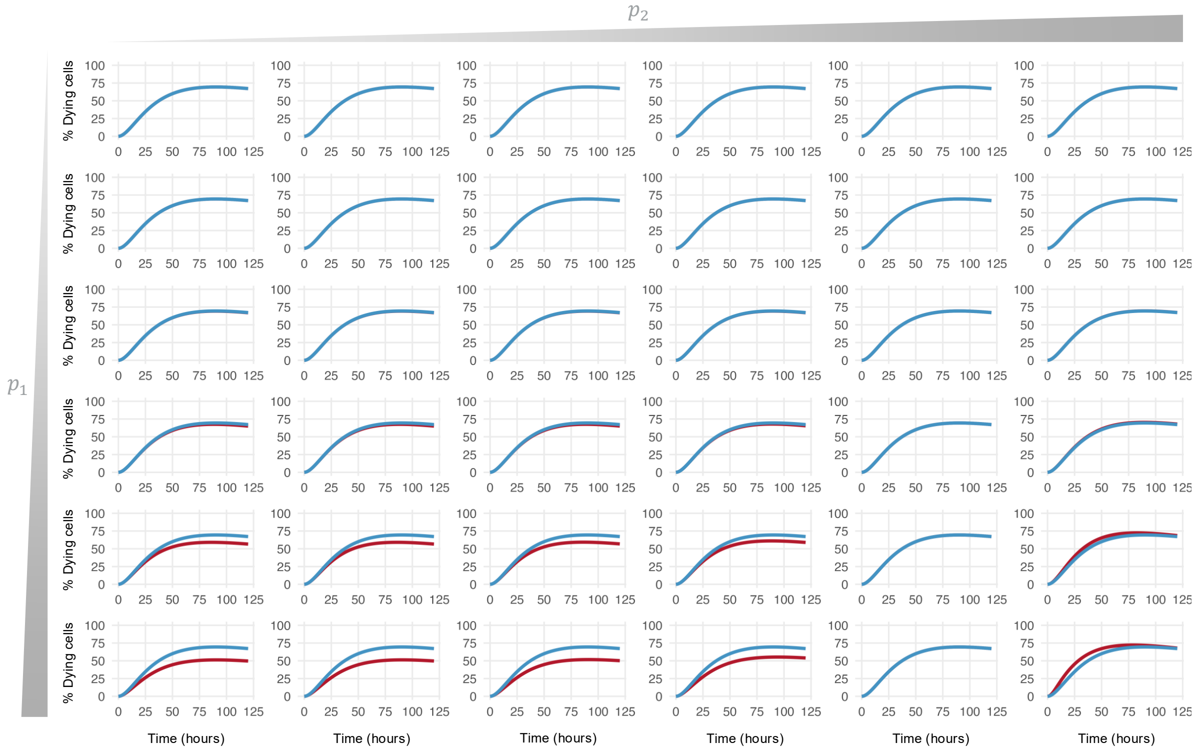


Figure S.5: **How differences in resistant-state dynamics affect treatment efficacy.** Simulations of the TNBC ODE model including drug-resistant states (SI–Eq. S.18) for $p_1 = 0, 0.001, 0.01, 0.1, 1, 10$ h$^{-1}$ and $p_2 = 0, 0.001, 0.01, 0.1, 1, 10$ h$^{-1}$, with $A = 0.0045$ h$^{-1}$ and $B = 0.0035$ h$^{-1}$ (red curves). The blue curve corresponds to the simulation of the TNBC model with $p_1 = p_2 = 0.5$ h$^{-1}$. The full set of parameter values is provided in SI–Section S.5.

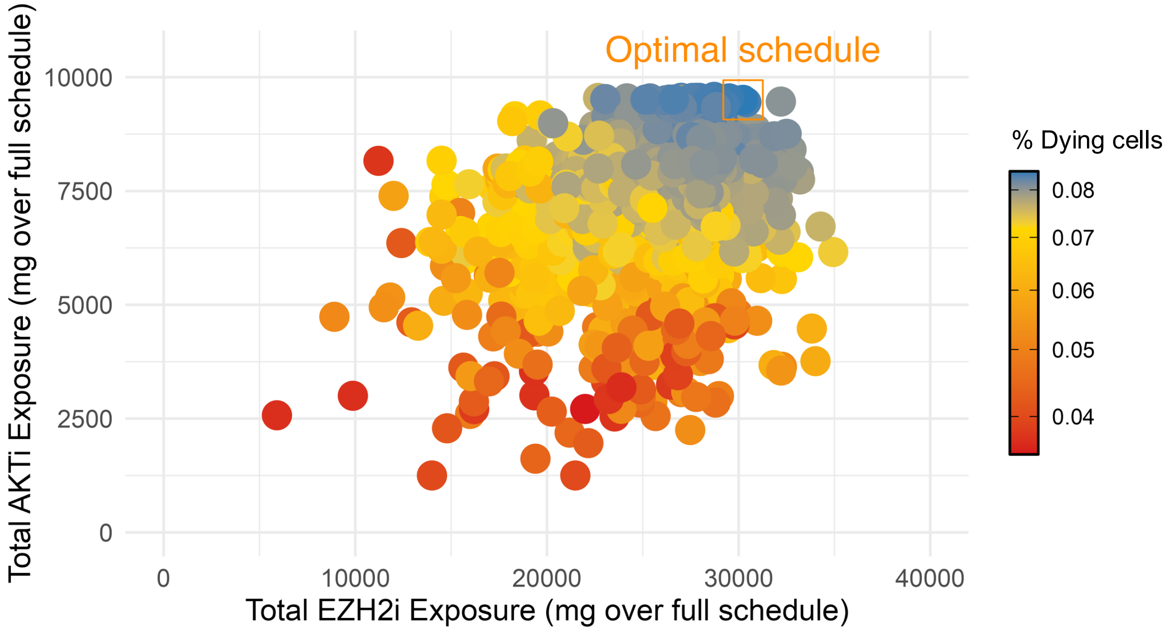


Figure S.6: **Treatment schedules evaluated by the genetic algorithm for regimens including capivasertib.** The genetic algorithm (GA) evaluates N = 4034 treatment schedules, shown according to their total tazemetostat (EZH2i) exposure and total capivasertib (AKTi) exposure across the entire treatment duration. The orange square marks the schedule identified by the GA as optimal. See Methods for details about GA algorithm implementation.

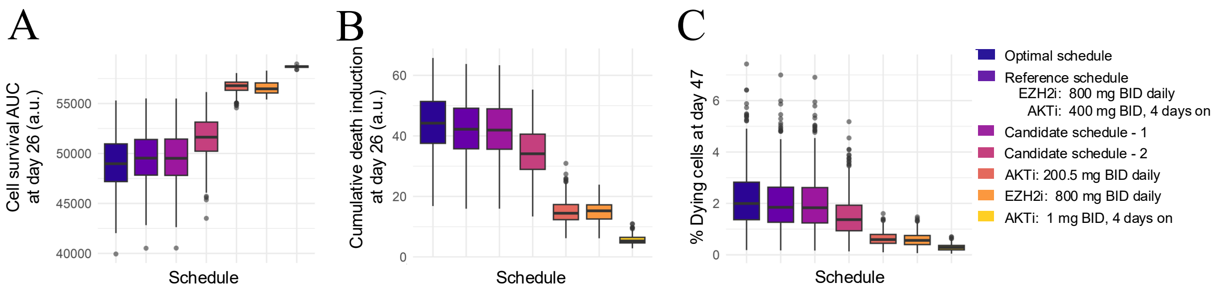


Figure S.7: **Additional evaluation metrics confirm the robustness of the optimized treatment schedule for the tazemetostat - capivasertib combination.** **A** Box plots of regimen efficacy quantified using the cell survival area under the curve (AUC) at the end of one treatment cycle (day 26) (see Methods). **B** Box plots of regimen efficacy quantified using cumulative death induction at the end of one treatment cycle (day 26) (see Methods). **C** Box plots of regimen efficacy quantified using % of dying cells after two treatment cycles (day 47). For all panels, boxes represent the middle 50% of the data, with the median shown as a horizontal line. Whiskers extend to the minimum and maximum values excluding outliers, which are shown as dots. Each regimen is represented with a different color and its results are based on N = 500 simulations, corresponding to 500 *in silico* patients. The optimal schedule and candidate schedules 1 and 2 are the same as introduced in Fig. 5C.

A

**Discretized optimal schedule**

| | Pretreat. (5 days) | Week 1 | Week 2 | Week 3 | Total |
|---|---|---|---|---|---|
| Tazemetostat (EZH2i) | 600 mg BID | 800 mg BID | 600 mg BID | 400 mg BID | 31200 mg |
| Ipatasertib (AKTi) | | 400 mg BID 7 days on 0 days off | 300 mg BID 4 days on 3 days off | 200 mg BID 4 days on 3 days off | 9600 mg |

**Discretized candidate schedule – 1**

| | Pretreat. (5 days) | Week 1 | Week 2 | Week 3 | Total |
|---|---|---|---|---|---|
| Tazemetostat (EZH2i) | 400 mg BID | 600 mg BID | 600 mg BID | 400 mg BID | 26400 mg |
| Ipatasertib (AKTi) | | 400 mg BID 6 days on 1 days off | 300 mg BID 3 days on 4 days off | 200 mg BID 1 days on 6 days off | 7000 mg |

**Discretized candidate schedule – 2**

| | Pretreat. (5 days) | Week 1 | Week 2 | Week 3 | Total |
|---|---|---|---|---|---|
| Tazemetostat (EZH2i) | 400 mg BID | 200 mg BID | 600 mg BID | 400 mg BID | 20800 mg |
| Ipatasertib (AKTi) | | 300 mg BID 5 days on 2 days off | 200 mg BID 1 days on 6 days off | 100 mg BID 4 days on 3 days off | 4200 mg |

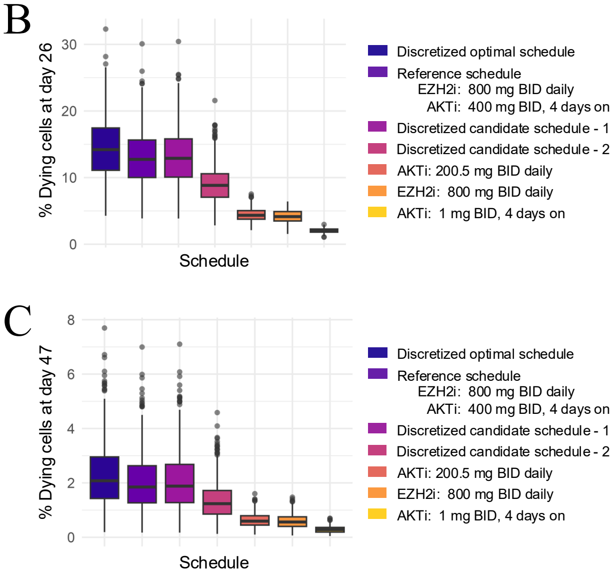


Figure S.8: **Evaluation of clinically feasible discretized treatment schedules for the tazemetostat - capivasertib combination.** **A** Summary of the optimal schedule and candidate schedules 1 and 2, introduced in Fig. 5C, converted into clinically feasible regimens by rounding each dose to the nearest allowable dose level. Allowed doses are 0, 200, 400, 600, and 800 mg BID for tazemetostat and 0, 100, 200, 300, and 400 mg BID for capviasertib. **B** Box plots of regimen efficacy quantified using % of dying cells after one treatment cycle (day 26) for the discretized schedules shown in panel A and the corresponding reference schedules. **C** Box plots of regimen efficacy quantified as % of dying cells after two treatment cycles (day 47) for the same regimens. For panels B and C, boxes represent the middle 50% of the data, with the median shown as a horizontal line. Whiskers extend to the minimum and maximum values excluding outliers, which are shown as dots. Each regimen is represented with a different color and its results are based on N = 500 simulations, corresponding to 500 *in silico* patients.

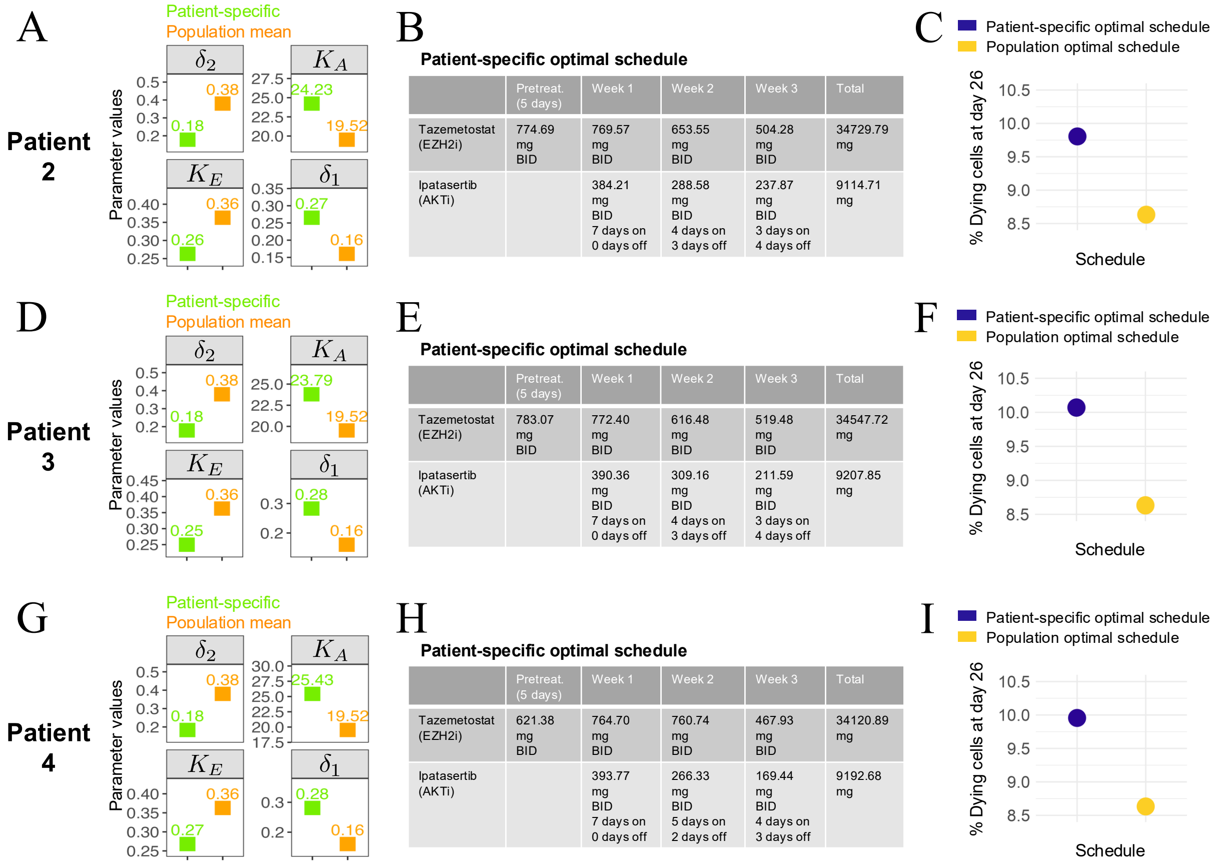


Patient 2 (B):

| | Pretreat. (5 days) | Week 1 | Week 2 | Week 3 | Total |
|---|---|---|---|---|---|
| Tazemetostat (EZH2i) | 774.69 mg BID | 769.57 mg BID | 653.55 mg BID | 504.28 mg BID | 34729.79 mg |
| Ipatasertib (AKTi) | | 384.21 mg BID 7 days on 0 days off | 288.58 mg BID 4 days on 3 days off | 237.87 mg BID 3 days on 4 days off | 9114.71 mg |

Patient 3 (E):

| | Pretreat. (5 days) | Week 1 | Week 2 | Week 3 | Total |
|---|---|---|---|---|---|
| Tazemetostat (EZH2i) | 783.07 mg BID | 772.40 mg BID | 616.48 mg BID | 519.48 mg BID | 34547.72 mg |
| Ipatasertib (AKTi) | | 390.36 mg BID 7 days on 0 days off | 309.16 mg BID 4 days on 3 days off | 211.59 mg BID 3 days on 4 days off | 9207.85 mg |

Patient 4 (H):

| | Pretreat. (5 days) | Week 1 | Week 2 | Week 3 | Total |
|---|---|---|---|---|---|
| Tazemetostat (EZH2i) | 621.38 mg BID | 764.70 mg BID | 760.74 mg BID | 467.93 mg BID | 34120.89 mg |
| Ipatasertib (AKTi) | | 393.77 mg BID 7 days on 0 days off | 266.33 mg BID 5 days on 2 days off | 169.44 mg BID 4 days on 3 days off | 9192.68 mg |

Figure S.9: **Application of the digital twin model to individualized treatment optimization for additional representative virtual patients. A** Comparison, for Patient 2, between the patient-specific parameter estimates obtained after calibration and the corresponding population-level mean parameter values, previously estimated from the population-level calibration (see SI-Section S.2), for the four patient-specific model parameters governing treatment response ($\delta_2$, $K_A$, $K_E$, and $\delta_1$). **B** Summary of the Patient 2 optimal treatment schedule. **C** Comparison of the predicted therapeutic response obtained using the Patient 2-specific optimal schedule and the population-level optimal schedule. The population-level optimal schedule is the same as the one introduced in Fig. 3E. **D,E,F** Same as panels A,B,C, but for Patient 3. **G,H,I** Same as panels A,B,C, but for Patient 4.

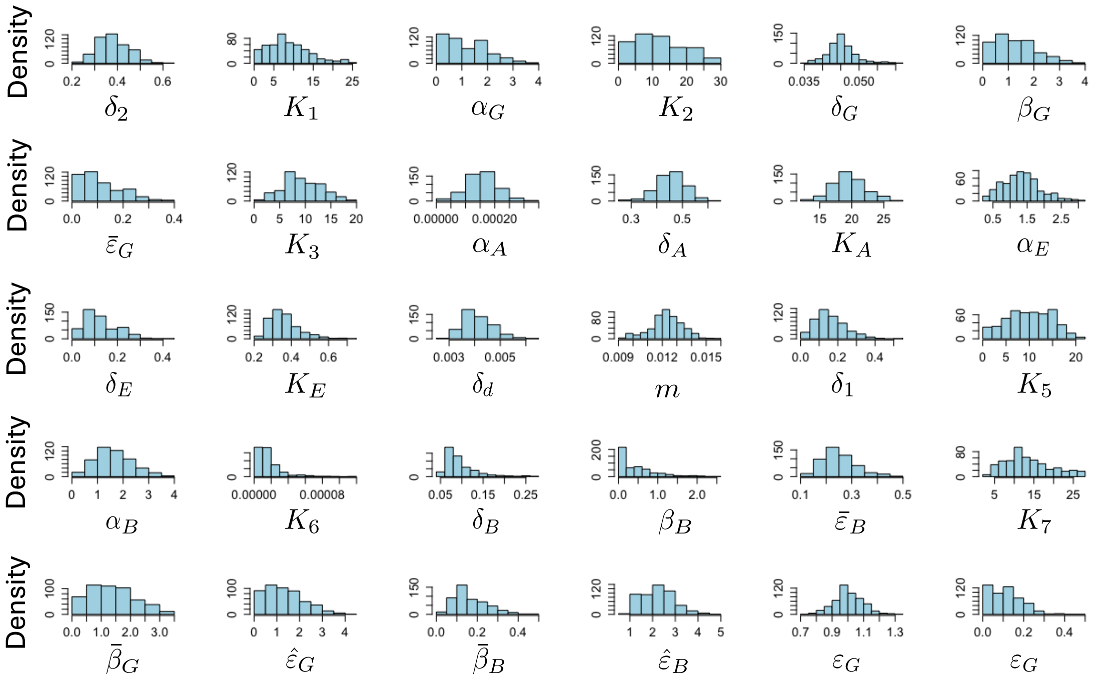


Figure S.10: **Posterior distributions of the parameters estimated in the epigenetic TNBC progression model.** Histograms showing the posterior distributions of the epigenetic TNBC progression model parameters obtained through Bayesian inference using the *in vitro* treatment response data in Fig. 2B (see Methods).

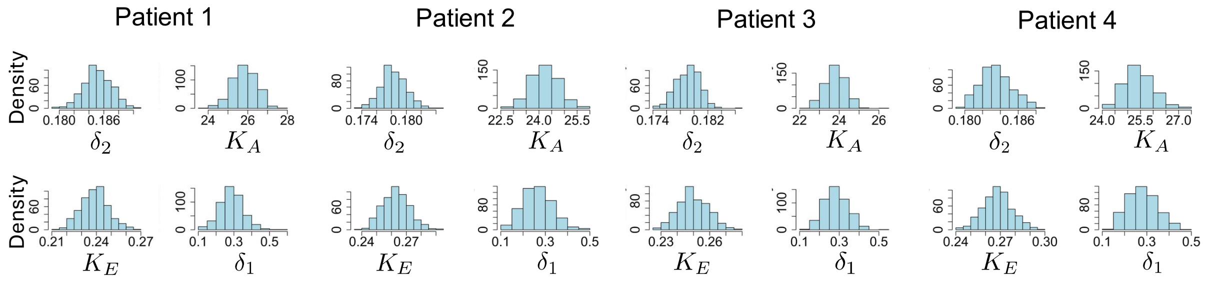


Figure S.11: **Posterior distributions of the patient-specific parameters estimated for four representative virtual patients.** Histograms showing the posterior distributions of the epigenetic TNBC progression model parameters obtained through Bayesian inference using the synthetic, patient-specific *in vitro* treatment response data of four representative virtual patients (see Methods). For each virtual patient, only the four patient-specific parameters ($\delta_2$, $K_A$, $K_E$, and $\delta_1$) were re-estimated, while the remaining model parameters were fixed to their population-level mean estimates (see Methods).
